# Functional constraints of *wtf* killer meiotic drivers

**DOI:** 10.1101/2024.08.27.609905

**Authors:** Ananya Nidamangala Srinivasa, Samuel Campbell, Shriram Venkatesan, Nicole L. Nuckolls, Jeffrey J. Lange, Randal Halfmann, Sarah E. Zanders

## Abstract

Killer meiotic drivers are selfish DNA loci that sabotage the gametes that do not inherit them from a driver+/driver- heterozygote. These drivers often employ toxic proteins that target essential cellular functions to cause the destruction of driver- gametes. Identifying the mechanisms of drivers can expand our understanding of infertility and reveal novel insights about the cellular functions targeted by drivers. In this work, we explore the molecular mechanisms underlying the *wtf* family of killer meiotic drivers found in fission yeasts.

Each *wtf* killer acts using a toxic Wtf^poison^ protein that can be neutralized by a corresponding Wtf^antidote^ protein. The *wtf* genes are rapidly evolving and extremely diverse. Here we found that self-assembly of Wtf^poison^ proteins is broadly conserved and associated with toxicity across the gene family, despite minimal amino acid conservation. In addition, we found the toxicity of Wtf^poison^ assemblies can be modulated by protein tags designed to increase or decrease the extent of the Wtf^poison^ assembly, implicating assembly size in toxicity. We also identified a conserved, critical role for the specific co-assembly of the Wtf^poison^ and Wtf^antidote^ proteins in promoting effective neutralization of Wtf^poison^ toxicity. Finally, we engineered *wtf* alleles that encode toxic Wtf^poison^ proteins that are not effectively neutralized by their corresponding Wtf^antidote^ proteins. The possibility of such self-destructive alleles reveals functional constraints on *wtf* evolution and suggests similar alleles could be cryptic contributors to infertility in fission yeast populations. As rapidly evolving killer meiotic drivers are widespread in eukaryotes, analogous self-killing drive alleles could contribute to sporadic infertility in many lineages.

**Author Summary:** Diploid organisms, such as humans, have two copies of most genes. Only one copy, however, is transmitted through gametes (e.g., sperm and egg) to any given offspring. Alternate copies of the same gene are expected to be equally represented in the gametes, resulting in random transmission to the next generation. However, some genes can "cheat" to be transmitted to more than half of the gametes, often at a cost to the host organism. Killer meiotic drivers are one such class of cheater genes that act by eliminating gametes lacking the driver. In this work, we studied the *wtf* family of killer meiotic drivers found in fission yeasts. Each *wtf* driver encodes a poison and an antidote protein to specifically kill gametes that do not inherit the driver.

Through analyzing a large suite of diverse natural and engineered mutant *wtf* genes, we identified multiple properties—such as poison self-assembly and poison-antidote co-assembly—that can constrain poison toxicity and antidote rescue. These constraints could influence the evolution of *wtf* genes. Additionally, we discovered several incompatible *wtf* poison-antidote pairs, demonstrating expanded potential for self-killing *wtf* alleles. Such alleles could potentially arise spontaneously in populations cause infertility.

## Introduction

Genomes often contain selfish DNA sequences that persist by promoting their own propagation into the next generation without providing an overall fitness benefit to the organism [1,2]. Killer meiotic drivers are one class of selfish sequences that act by preferentially destroying gametes that do not inherit the driver from a heterozygote [3]. This gamete destruction leads to biased, or sometimes, complete transmission of the driver+ genotype from driver+/driver- heterozygotes.

Killer meiotic drive systems generally decrease the fitness of the organism carrying the driver, both through their killing activities and through indirect mechanisms [4].

Distinct killer meiotic drive systems have repeatedly evolved in eukaryotes and drivers found in distinct species are generally not homologous [3,5–12]. Despite this, killer meiotic drivers fall into a limited number of mechanistic classes with shared themes [3,13,14]. Additionally, unrelated killer meiotic drivers have recurrently exploited conserved facets of cell physiology.

This has enabled study of the killing and/or rescue activities of drive proteins outside of their endogenous species [5,11,15,16]. Overall, drivers represent unique tools that can be used to discover novel, unexpected insights into the exploited biological processes.

The *wtf* genes are a family of extremely diverse, rapidly evolving killer meiotic drivers found in multiple copies in most *Schizosaccharomyces* (fission yeast) species [17–22]. The *wtf* driver genes each produce a Wtf^poison^ and a Wtf^antidote^ protein using distinct transcripts with overlapping coding sequences (Fig 1A) [17,22,23]. The amino acid sequences of the two proteins are largely identical, except the Wtf^antidote^ proteins have an additional N-terminal domain of about 45 amino acids not found in the Wtf^poison^ (Fig 1A). All four developing spores (products of meiosis) are exposed to the Wtf^poison^, but only spores that inherit a compatible Wtf^antidote^ can neutralize the poison and survive [22].

**Figure 1.**
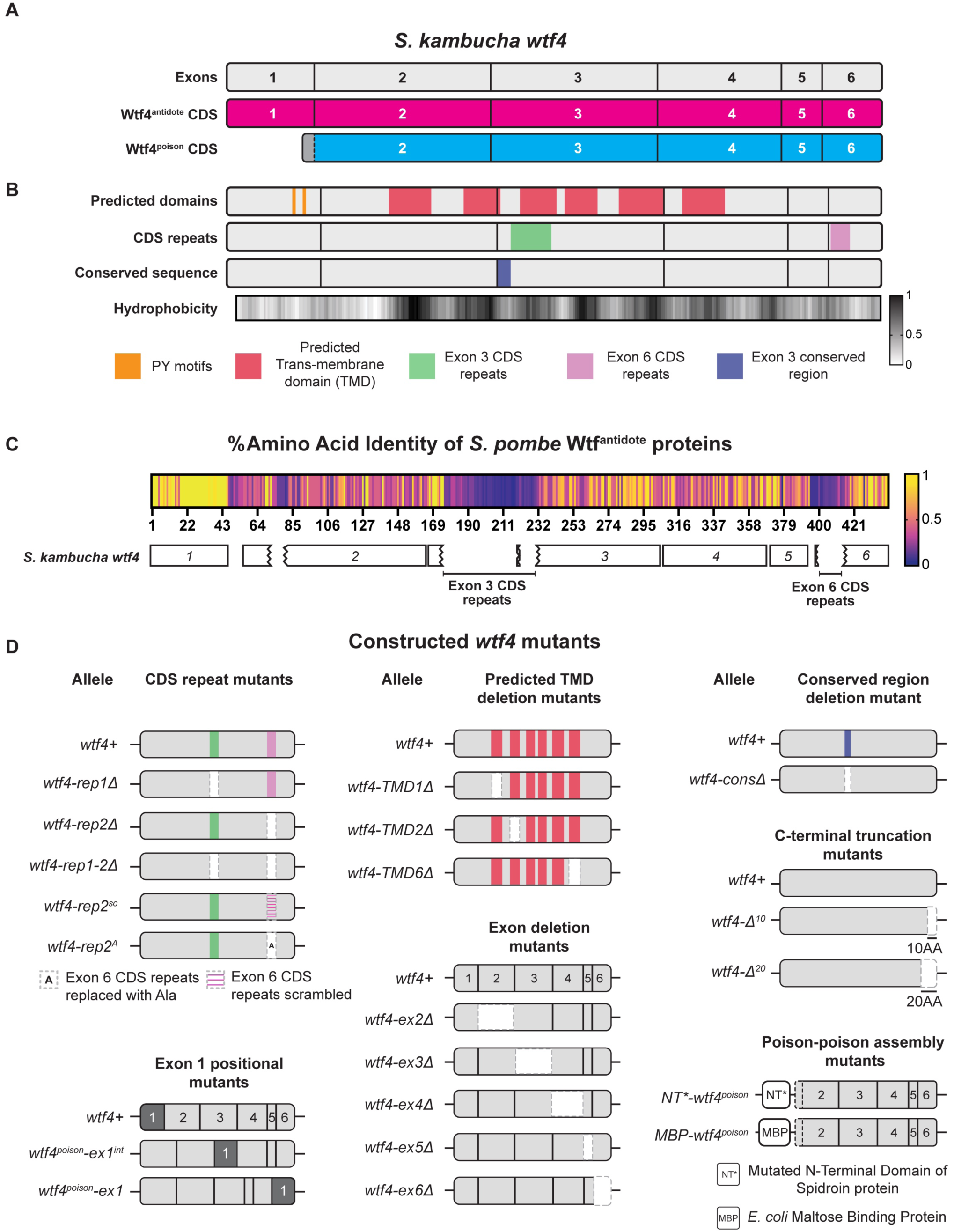
Features of *wtf4* and mutant alleles. **A.** A cartoon of *S. kambucha wtf4* coding sequence (CDS). Wtf4^antidote^ coding sequence is shown in magenta, which includes exons 1-6. The Wtf4^poison^ coding sequence is shown in cyan, which includes 21 base pairs from intron 1 (in grey), and exons 2-6. **B.** Features of Wtf4^antidote^ and Wtf4^poison^ proteins. Row 1 shows predicted secondary structure domains and functional motifs, including PY motifs (in mustard) and predicted transmembrane domains (in red). Row 2 shows the coding sequence repeats in exon 3 and exon 6. Row 3 highlights a well-conserved region 9 amino acids long. Row 4 is the normalized hydrophobicity of the Wtf4 proteins from ProtScale, with the Kyle and Doolittle Hydropathy scale [81]. The higher the number on the scale, the higher the hydrophobicity of the amino acid. See S5 and S6 Tables for detailed descriptions. **C.** Percentage amino acid identity of Wtf^antidotes^ from 33 *wtf* driver genes from four isolates of *S. pombe* [20]. The antidote sequences were aligned using Geneious Prime (2023.0.4) and the percentage amino acid identity is depicted as a heatmap, with yellow being 100% identity. *S. kambucha wtf4* CDS is shown below for comparison, with the exons labeled corresponding to the consensus. Labeled areas within exons 3 and 6, where identity is low, represent the expansion and contraction of the coding sequence repeats of different *wtf* genes. **D.** Cartoon of *wtf4* mutants constructed in this study. Each mutant category was constructed based on a specific feature mentioned above. The categories are depicted with a wild type *wtf4* allele at the top. See S1 Table for a comprehensive overview of the alleles and their phenotypes.

Little is known about the mechanism of toxicity of the Wtf^poison^ proteins. The general mechanism used by Wtf^antidote^ proteins is better understood. The antidote-specific N-terminal domain is the most conserved region and includes targeting motifs (PY motifs) that are recognized by Rsp5/NEDD4 ubiquitin ligases which route the protein through the trans Golgi network to endosomes and, ultimately, to the vacuole (fungal lysosome; Fig 1B) [15,24]. When both proteins are present, the Wtf^antidote^ co-assembles with the Wtf^poison^, and the Wtf^antidote^ thereby traffics the Wtf^poison^ to the vacuole [15,24]. This mechanism is similar to a non-homologous drive system recently described in rice, where an antidote protein rescues the gametes that inherit the drive locus from a toxic poison protein by co-trafficking the poison to the autophagosome [5].

In general, the poison protein encoded by one *wtf* gene is not compatible with (i.e., neutralized by) the antidotes of widely diverged *wtf* genes [17,19,25]. However, Wtf^poison^ proteins can be neutralized by Wtf^antidote^ proteins encoded at different loci if the sequences are identical or highly similar (outside of the antidote-specific N-terminal domain; Fig 1A) [17,22,24,26]. Although the examples are limited, similarity at the C-termini of the Wtf poison and antidote proteins may be particularly important for compatibility [15,25,26]. For example, the *wtf18-2* allele encodes an antidote protein that neutralizes the poison produced by the *wtf13* driver [26]. The Wtf18-2^antidote^ and Wtf13^antidote^ proteins are highly similar overall (82% amino acid identity) and identical at their C-termini. Interestingly, the antidote encoded by the reference allele of *wtf18* is not identical to Wtf13^poison^ at the C-terminus and does not neutralize it. In addition, swapping one amino acid for two different amino acids in the C-terminus of Wtf18-2^antidote^ (D366NN mutation) to make it more like the reference Wtf18^antidote^ abolishes the protein’s ability to neutralize Wtf13^poison^ [26]. Still, the rules governing poison and antidote compatibility are largely unknown.

While it is clear that a functional *wtf* driver kills about half the spores produced by *wtf*+/*wtf*- heterozygotes, the full impacts of *wtf* gene evolution on the fitness of populations is not clear. Because the *wtf* genes encode the poison and antidote proteins on overlapping coding sequences (Fig 1A), novel poisons can emerge simultaneously with their corresponding antidotes via mutation. This has been observed with a limited number of engineered *wtf* alleles, and one can observe evidence of this divergence in the diversity of extant *wtf* alleles in natural populations [15,19–21,26]. The natural alleles, however, represent a selected population and thus provide a biased sample of the novel *wtf* alleles generated by mutation and recombination. Novel alleles that generate functional drivers are predicted to be favored by the self-selection enabled by drive and be over-represented in natural populations. Conversely, alleles that generate a toxic poison without a compatible antidote would be expected to be under- represented in natural populations due to infertility caused by self-killing.

Major questions remain about the mechanism(s) of Wtf^poison^ toxicity and about the rules of Wtf^poison^ and Wtf^antidote^ compatibility. Our working model is that the toxicity of Wtf^poison^ proteins is tied to their self-assembly [15]. This assembly, however, has only been conclusively demonstrated for the Wtf4 proteins, encoded by the *wtf4* gene from the isolate of *S. pombe* known as *kambucha*. For Wtf^antidote^ function, a working model is that a similar homotypic assembly with the Wtf^poison^ is required to establish a physical connection between the proteins so that the poison is shuttled to the vacuole along with a ubiquitinated antidote [15,24]. The ubiquitination and trafficking of the Wtf antidotes to the vacuole has been demonstrated to be conserved between widely diverged Wtf^antidote^ proteins [19,24]. However, it is not clear if antidote co-assembly with a poison has a functional role in poison neutralization, or if it merely provides a physical link between the proteins to facilitate co-trafficking. It is also unclear if changes in the coding sequence shared by the poison and antidote proteins of a given *wtf* can generate an incompatible set of proteins, or if poisons will always be neutralized by sequence-matched antidotes.

To test these models and to better understand the amino acid sequences that support the Wtf protein functions, we analyzed a panel of natural and engineered Wtf proteins. We found that both well-conserved and poorly conserved amino acid sequences can contribute to protein function. Our analyses revealed broad conservation of Wtf^poison^ self-assembly and suggest that assembly size can affect Wtf^poison^ toxicity, analogous to several other self-assembling toxic proteins [27–29]. This strongly implicates self-assembly as a critical parameter in Wtf^poison^ toxicity. In addition, we found that specific co-assembly with the Wtf^antidote^ is required for efficient Wtf^poison^ neutralization via trafficking to the vacuole. Finally, our analyses identified multiple *wtf* alleles that generate poison proteins that are not efficiently neutralized by their corresponding antidotes. Such alleles are self-destructive and could contribute to sporadic infertility. This work refines our understanding of the functional constraints of Wtf proteins, with important evolutionary implications. More broadly, our observations offer insight into how functional conservation can be maintained despite extreme amino acid sequence divergence and extends our understanding of the limits of protein assembly trafficking mechanisms.

## Results

### Wtf^poison^ protein self-assembly is broadly conserved

To test the models of Wtf protein functions and to understand how these functions could be supported by extremely diverse protein sequences (e.g., <20% amino acid identity), we generated a large panel of *wtf* variants that represent over 100 million years of divergence [19]. We reasoned that functionally important features of the proteins would be conserved in functional variants (i.e. toxic Wtf^poison^ proteins), despite minimal amino acid conservation (Fig 1C). Twelve of the variants we assayed are wild-type or mutant alleles of *wtf* genes found in *S. octosporus*, *S. osmophilus* or *S. cryophilus* (S1 Fig). One of these genes, *S. octosporus wtf25*, has been previously characterized and found to encode a functional meiotic driver in its endogenous species ([19]; S1 Table). Four additional genes (*S. cryophilus wtf1*, *S. osmophilus wtf41*, and *S. octosporus wtf61*) have been shown to encode functional Wtf^poison^ and Wtf^antidote^ proteins in an ectopic *Saccharomyces cerevisiae* assay system ([19]; S1 Table). The *S. cerevisiae* system used to study Wtf proteins is well established and reflects phenotypes observed in the endogenous fission yeast species [15,19,24]. Finally, we assayed a panel of new mutant alleles of the previously characterized *wtf4* gene from the *S. kambucha* isolate of *S. pombe* [15,22,23] (Fig 1D).

Our initial model, based on Wtf4^poison^, was that Wtf^poison^ toxicity is linked to the ability of a Wtf^poison^ to self-assemble [15]. To test this idea, we assayed if other functional Wtf^poison^ proteins besides wild-type Wtf4^poison^ also self-assemble. To do this, we used the Distributed Amphifluoric FRET (DAmFRET) approach, in which proteins are tagged with the photoconvertible mEos3.1 fluorophore (referred to as mEos hereafter) and expressed in *S. cerevisiae*. A uniform fraction of the green mEos population is photoconverted to its red form, and protein-protein interactions are then detected as FRET (Fluorescence Resonance Energy Transfer) between the green and red fluorophores within single cells. Monomeric mEos exhibits negligible FRET and is used as a negative control (Fig 2A and 2B) [15,30].

**Figure 2.**
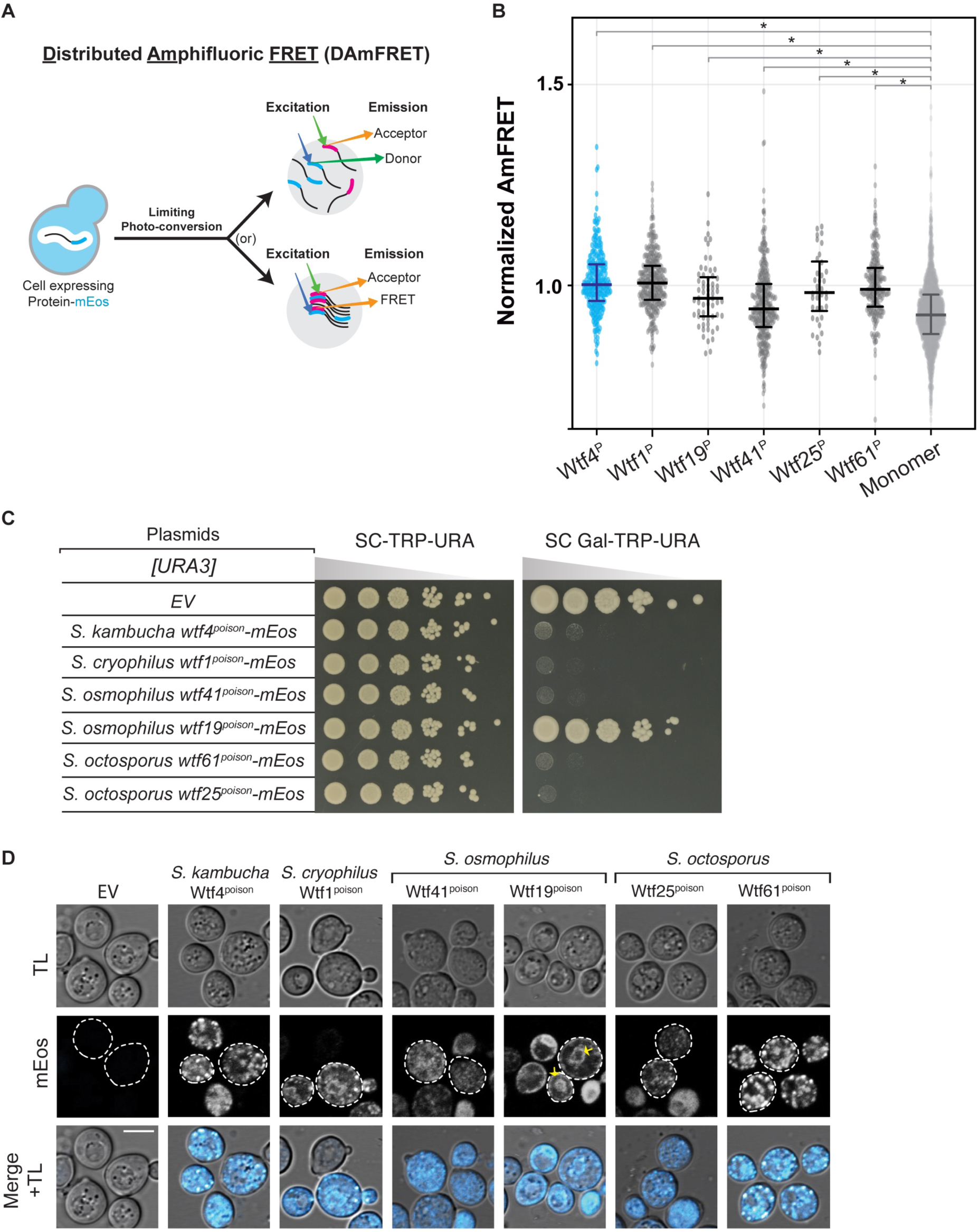
Extremely diverged toxic Wtf^poison^ proteins exhibit similar self-assembly and intracellular localization. **A**. Cartoon illustrating the Distributed Amphifluoric FRET (DAmFRET) assay (modified from [30]). AmFRET values for individual cells were calculated by dividing the acceptor fluorescence by the FRET fluorescence, both measured via cytometry. **B.** Combined AmFRET values for three technical replicates of the specified Wtf^poison^-mEos proteins and monomer-mEos (negative control). X^P^ represents the WtfX poison proteins tested here. The median is indicated with a solid line and the bars represent the interquartile range. For easier comparison, the values were normalized so that Wtf4^poison^ had a median of 1 in each experiment. The data shown here do not include outliers. See S2 Data for the complete dataset and p-values. Statistical significance: *p<0.01; t-tests with Bonferroni correction. **C.** A spot assay of cells serially diluted and plated on SC-TRP-URA and SC Gal-TRP-URA plates. Each strain carries an empty [*TRP1*] plasmid, and either an empty [*URA3*] plasmid (EV) or the indicated *wtf^poison^-mEos* allele under the control of a galactose-inducible promoter. The plates were grown at 30^·^C for 3 days. **D.** Representative images of the same strains depicted in C were induced in galactose media for 4 hours at 30^·^C to express the indicated mEos-tagged proteins. The images are not at the same brightness and contrast settings to clearly show localization of tagged proteins. Yellow arrows indicate endoplasmic reticulum-like localization. TL is transmitted light, and the scale bar is 4 µm.

We first tagged four toxic (*S. cryophilus* Wtf1^poison^, *S. osmophilus* Wtf41^poison^, and *S. octosporus* Wtf25^poison^ and Wtf61^poison^) and one non-toxic (*S. osmophilus* Wtf19^poison^) Wtf^poison^ protein at the C-terminus with mEos. The tagged proteins retained their toxic/non-toxic phenotypes when expressed in *S. cerevisiae* using a galactose-inducible expression system (Fig 2C) [19]. The assayed proteins all share less than 20% amino acid identity with Wtf4^poison^, which we use as a positive control (S1B Fig). We found that all five newly tested Wtf^poison^-mEos proteins, including the nontoxic Wtf19^poison^-mEos, exhibited self-assembly (Fig 2B). It is important to note, however, that the data reported for Wtf25^poison^-mEos are limited and we could not reliably assay that protein via DAmFRET. This is because only living cells are considered in DAmFRET and Wtf25^poison^-mEos kills cells rapidly at low expression (e.g. they die faster than cells expressing Wtf4^poison^)_._

We also assayed DAmFRET in partial deletion alleles of *wtf4^poison^-mEos* (S2 and S3 Figs). Some of these alleles are described in more detail below, but for this section, we focused on exploring the potential connection between assembly and toxicity. The two toxic mutant Wtf4^poison^ proteins tested, Wtf4-ex5Δ^poison^-mEos (S2A and S2D-F Fig) and Wtf4Δ^20-poison^-mEos (S3A-E Fig) both exhibited self-assembly. Of these two proteins, Wtf4Δ^20-poison^-mEos is more like wild-type Wtf4^poison^-mEos, in both toxicity and degree of assembly (S2D-F and S3B-E Figs).

Most non-toxic Wtf4 mutant proteins (Wtf4-TMD1Δ^poison^-mEos, Wtf4-ex3Δ^poison^-mEos, Wtf4- ex4Δ^poison^-mEos, Wtf4-ex6Δ^poison^-mEos and Wtf4-consΔ^poison^-mEos) exhibited reduced assembly compared to the wild-type (S2A-E and S3A-D Figs). We did, however, find one exceptional mutant protein, Wtf4Δ^10-poison^-mEos. This protein was not toxic but showed similar assembly to the wild-type protein (S3A-D Fig). We could not reliably assay the assembly of some alleles (Wtf4-TMD2Δ^poison^-mEos, Wtf4-TMD6Δ^poison^-mEos and Wtf4-ex2Δ^poison^-mEos) due to low fluorescence signal (S2A, S2B and S4 Figs).

These results show that Wtf^poison^ self-assembly is broadly conserved (over 100 million years) and can be supported by a wide range of amino acid sequences (e.g., proteins sharing <20% amino acid identity, S1B Fig). Given the broad conservation of self-assembly, that all the toxic alleles exhibited assembly, and that mutant alleles with reduced toxicity also exhibited reduced assembly, our results strongly support an association between self-assembly and toxicity.

Importantly, our results also demonstrate that self-assembly alone is insufficient for Wtf^poison^ toxicity, as we found several Wtf^poison^ proteins (Wtf19-mEos and Wtf4Δ^10-poison^-mEos) that assemble but are not toxic.

### Deletion alleles demonstrate that both conserved and non-conserved regions contribute to Wtf4^poison^ self-assembly and toxicity

Our deletion mutants in *wtf4^poison^* encompass both non-conserved and conserved sequences allowing us to test the functional importance of both sequence types (Fig 1; [19–21]). Within *S. pombe* Wtf proteins, there is one well conserved 29 base pair segment found at the beginning of exon 3 (Fig 1B; [20]). The Wtf4-consΔ^poison^-mEos protein lacks this domain and has disrupted self-assembly and is non-toxic, indicating the conserved sequence has a functional role in both Wtf4^poison^ assembly and toxicity (S2C-E Fig).

We similarly found that several regions that are poorly conserved in Wtf proteins are functionally important in Wtf4^poison^. For example, exon 4 is not conserved within functional *wtf* genes, even within *S. pombe* [19, 20, 25]. Still, our results demonstrate exon 4 is functionally important in *wtf4* as the Wtf4-ex4Δ^poison^-mEos protein has disrupted self-assembly and is not toxic (S2A, S2D and S2E Fig). An additional variable feature of *wtf* genes is the number of predicted transmembrane domains, which varies between four and eight within functional Wtf proteins (S2 Table). The *wtf4* gene has 6 predicted transmembrane domains and we assayed individual deletions of three (Wtf4-TMD1Δ^poison^-mEos, Wtf4-TMD2Δ^poison^-mEos and Wtf4-TMD6Δ^poison^- mEos). All three transmembrane deletions disrupted toxicity, indicating they are also functionally important (S2B, S2D and S2E Fig).

An additional set of deletion mutants we queried in a poorly conserved region followed up on an observation by Hu et al [17] in one of the inaugural papers describing *wtf* drivers. In that work, Hu et al [17] described 10 amino acid C-terminal truncations of two *wtf* genes known as *cw9* and *cw27*. These mutant alleles both exhibited disrupted Wtf^poison^, but not Wtf^antidote^ activity, implicating the C-terminus in Wtf^poison^ function. These proteins share 62% amino acid identity in the C-terminal exon 6 with each other and, at most, 70% with Wtf4. We tested if a similar 10 amino acid truncation of *wtf4* would specifically disrupt Wtf4^poison^ activity. This mutation gave us the exceptional *wtf4-Δ^10^* allele described above that encodes a non-toxic Wtf4Δ^10-poison^, that surprisingly exhibits wild-type self-assembly (S3A-D Fig). To expand our analyses beyond *wtf4*, we also made 10 amino acid C-terminal truncations of *S. cryophilus* Wtf1*, S. osmophilus* Wtf41*, S. octosporus* Wtf61 and Wtf25 poison. These proteins are all significantly smaller than Wtf4 (S1C Fig). Surprisingly, we found that these additional truncated proteins all retained toxicity (S3F and S3G Fig) and could be rescued by the corresponding antidotes (S5 Fig) indicating the C-termini are not universally important for Wtf^poison^ function.

We extended our deletion analyses of the C-terminus of Wtf4 poison and were surprised to find that deleting ten more amino acids than the ten missing in the non-toxic Wtf4Δ^10-poison^ protein restored toxicity. Specifically, the Wtf4Δ^20-poison^ protein exhibited robust self-assembly and near wild-type levels of toxicity (S3A-D Fig). A larger deletion of 29 amino acids, Wtf4ex6Δ^poison^, showed reduced self-assembly and was not toxic (S3A-D Fig).

Together, our results show that even non-conserved sequences can have context-dependent importance within a Wtf^poison^ protein. Specifically, a feature (e.g. the residues encoded in exon 4, or the last 10 amino acids of Wtf4^poison^) can be functionally important in one Wtf^poison^ protein but be missing or dispensable in another. This, combined with the lack of conservation within Wtf proteins, suggests that the contextually important amino acids (like the last 10 of Wtf4^poison^) do not have a specific function, but can rather contribute to the overall properties of the protein.

Changes in these contextually important regions can be complemented by changes elsewhere in the protein.

### Intracellular localization of mutant Wtf^poison^ proteins is correlated with toxicity

We identified self-assembly as one feature shared by functional Wtf^poison^ proteins and suspected that intracellular localization could be another, since we previously found that the toxic *S. octosporus* Wtf25^poison^-mCherry exhibited similar localization to that of toxic Wtf4^poison^ [19].

Specifically, both poisons show small puncta broadly distributed in the cytoplasm of *S. cerevisiae* cells, with minor localization to what appears to be the endoplasmic reticulum (ER; [15,19]). All functionally confirmed Wtf driver proteins have multiple predicted transmembrane domains (S2 and S3 Tables) and hence, this localization pattern may reflect the poison being trafficked from the ER through the secretory pathway, through the Golgi and trans-Golgi network [24].

To test the hypothesis that a link exists between the intracellular distribution and toxicity of Wtf^poison^ proteins, we imaged the wild-type Wtf^poison^-mEos proteins described above in *S. cerevisiae* cells. The localization of the *S. octosporus* Wtf25^poison^-mEos was similar to the previously described Wtf25^poison^-mCherry described above (Fig 2D). The additional toxic Wtf^poison^ proteins from *S. octosporus*, *S. osmophilus* and *S. cryophilus*, also exhibited dispersed puncta, like the toxic Wtf4^poison^-mEos control cells (Fig 2D). The nontoxic *S. osmophilus* Wtf19^poison^- mEos, however, showed a distinct localization pattern with strong signal enrichment in what appears to be the ER (Fig 2D). These data are consistent with our hypothesis that there is a link between intracellular distribution and toxicity with toxic Wtf^poison^ proteins exhibiting distributed puncta.

To further test our hypothesis, we also assayed the localization of the mutant Wtf4^poison^ proteins mentioned above (S2F and S3E Figs). The most toxic mutant protein, Wtf4Δ^20poison^-mEos, localized in distributed puncta like wild-type (S3A-C and S3E Fig). The less toxic Wtf4- ex5Δ^poison^-mEos protein also formed some distributed puncta, but also exhibited more ER-like localization than wild-type (S2A, S2D and S2F Fig). Most of the non-toxic mutant Wtf4^poison^ proteins (Wtf4-TMD1Δ^poison^-mEos, Wtf4-TMD2Δ^poison^-mEos, Wtf4-ex2Δ^poison^-mEos, Wtf4- ex3Δ^poison^-mEos, Wtf4-ex4Δ^poison^-mEos, and Wtf4-consΔ^poison^-mEos) were less dispersed and exhibited a largely ER-like localization, similar to the non-toxic *S. osmophilus* Wtf19^poison^-mEos (Fig 2C-D and S2A-D and S2F Fig). One additional non-toxic mutant, Wtf4-TMD6Δ^poison^-mEos, localized to the vacuole, reminiscent of Wtf^antidote^ protein localization (S2B, S2D and S2F Fig). The non-toxic Wtf4-ex6Δ^poison^-mEos had a unique localization pattern that combined a diffuse distributed signal with some larger protein assemblies (S3A-C and S3E Fig). Finally, the non-toxic Wtf4Δ^10-poison^-mEos localization was indistinguishable from wild-type Wtf4^poison^-mEos (S3A- C and S3E Fig).

In summary, all the toxic Wtf^poison^ proteins show a distributed puncta localization pattern, like Wtf4^poison^. All except one non-toxic Wtf^poison^ protein showed a distinct localization pattern from Wtf4^poison^, most often with enhanced ER-like localization. These results parallel our DAmFRET analyses where all toxic Wtf^poison^ proteins assemble and all the nontoxic proteins, except one, show reduced or lack of assembly. In both cases, the exceptional protein is Wtf4Δ^10-poison^, which is non-toxic, but shows wild-type assembly and localization in cells. Despite the exception, our results show that Wtf^poison^ self-assembly (assayed by DAmFRET) combined with cellular localization patterns are good predictors of Wtf^poison^ protein toxicity (Fig 7A).

### Altering assembly properties of Wtf^poison^ proteins affects toxicity

To further test the model that the toxicity of Wtf^poison^ proteins is tied to their self-assembly, we sought to alter the assembly properties of Wtf4^poison^ using tags. We first aimed to increase the overall size of the Wtf4^poison^ assemblies. To do this, we employed a recently described tool to oligomerize mEos3-fused proteins in trans. Specifically, we expressed a fusion protein consisting of the human ferritin heavy chain protein (FTH1) and a nanobody that recognizes mEos (mEosNB) [31]. The FTH1 domain self assembles to form a 24-mer core and can form supramolecular clusters when fused to self-assembling proteins [32,33]. In our assay, we expressed Wtf4^poison^-mEos from a galactose inducible promoter and the mEosNB-FTH1 from a doxycycline repressible promoter (Fig 3A and 3B). As expected, we found that Wtf4^poison^-mEos formed fewer and larger puncta inside cells in the presence of mEosNB-FTH1, consistent with the mEosNB-FTH1 complexes bringing Wtf^poison^-mEos assemblies into supramolecular clusters (i.e., -Dox panels, Fig 3C). Expression of the mEosNB-FTH1 alone had no effect on yeast growth (Fig 3D, panel (i)). While expression of the Wtf4^poison^-mEos alone was toxic, co-expression of the mEosNB-FTH1 and Wtf4^poison^-mEos markedly suppressed the toxicity of the Wtf4^poison^-mEos (Fig 3D, compare panel (ii) to panel (iii)). We also observed the same suppression of toxicity with four additional toxic Wtf^poison^-mEos proteins in the presence of the mEosNB-FTH1, indicating the effect was not specific to Wtf4^poison^ (Fig 3C and 3D). These results indicate that these supramolecular Wtf^poison^ assemblies are non-toxic. This effect could be due to altered intracellular localization and/or the reduced exposed surface area of the Wtf^poison^ assemblies.

**Figure 3.**
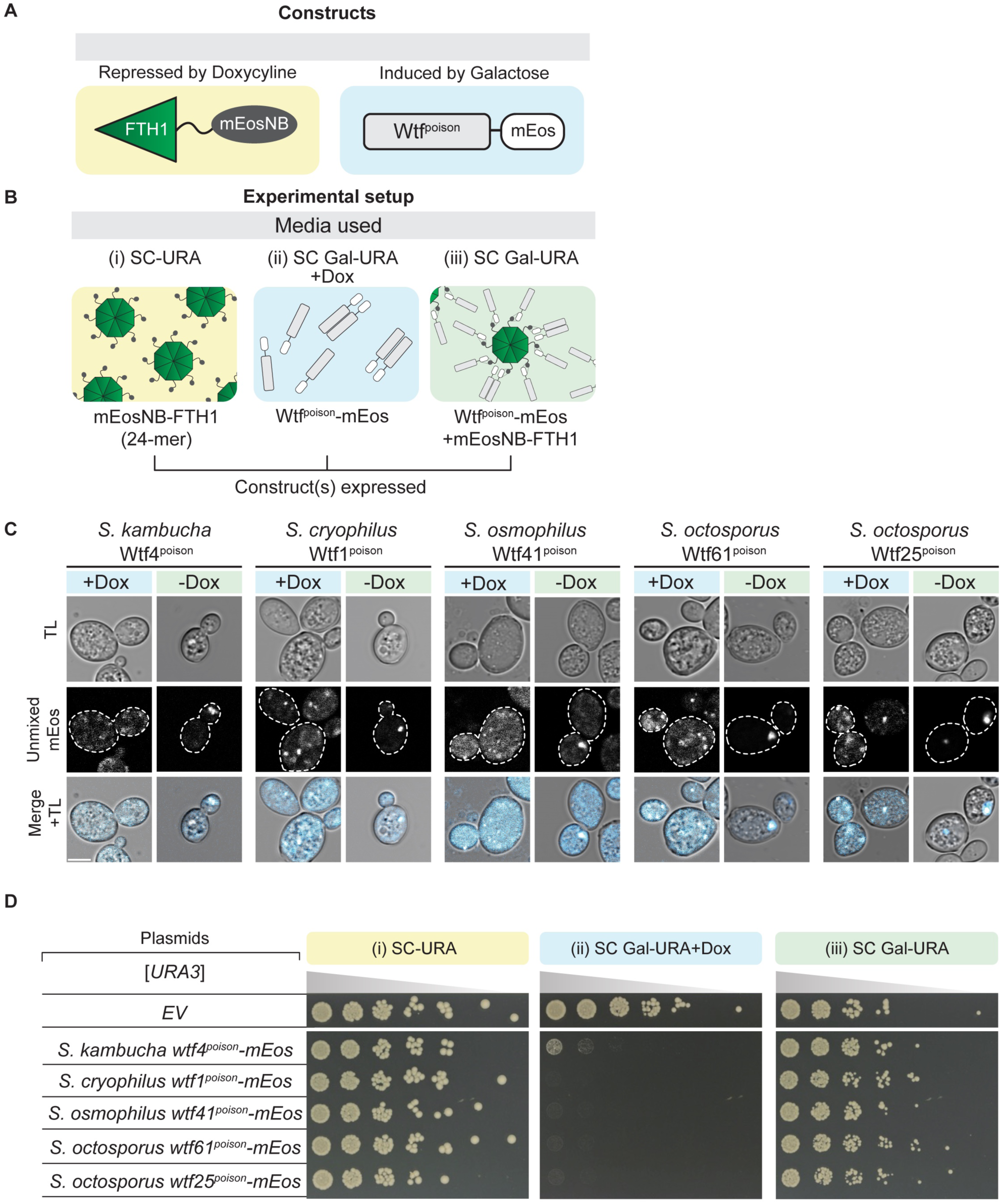
Increasing poison assembly with tags suppresses Wtf^poison^ toxicity. **A.** A cartoon of the constructs used in this experiment (C-D). The mEosNB-FTH1 construct was integrated into the genome and is under the control of a doxycycline-repressible promoter. mEosNB is a nanobody that binds mEos. The *wtf^poison^* alleles are carried on a [*URA3*] plasmid and are under the control of a galactose-inducible promoter. **B.** Cartoon of the constructs expressed on each medium used: SC-Ura, SC Galactose-Ura + 40 mg/L doxycycline, and SC Galactose-Ura. FTH1 is a 24-mer but is depicted as an 8-mer core. **C.** Representative images of cells induced with galactose media (with or without 40mg/L Doxycycline) for 4 hours at 30^·^C to express the indicated Wtf^poison^mEos proteins. The cells induced without Doxycycline (-Dox) also express mEosNB-FTH1. This strain background exhibits high autofluorescence, so we spectrally unmixed the signal to remove autofluorescence (See Methods). The images are not at the same brightness and contrast settings to clearly show localization of tagged proteins. TL is transmitted light, and the scale bar is 4 µm. **D.** A spot assay of cells carrying the constructs illustrated in A-C serially diluted on SC-Ura, SC Galactose-Ura + 40mg/L Doxycycline and SC Galactose-Ura plates and grown at 30^·^C for 3 days. Each strain carries either an empty [*URA3*] plasmid (EV) or the indicated *wtf^poison^-mEos* allele. These media induce the expression of mEosNB-FTH1 (i), the indicated Wtf^poison^-mEos protein (ii), or both (iii), respectively. The horizontal break in the image of each plate is due to rearrangements of the images to facilitate easy comparison. All strains within a panel were grown on the same plates (i.e., one SC-TRP-URA or SC Gal-TRP-URA or SC Gal-TRP-URA + 40mg/L Doxycycline plate).

We next attempted to increase the solubility Wtf4^poison^ assemblies using a modified N-terminal domain (NT*) tag derived from spidroins, a principal component of spider silk [34]. The NT* tag can decrease protein aggregation, which is the role of the native domain in spiders [35–39]. We added NT* to the N-terminus of the Wtf4^poison^-mEos to generate NT*-Wtf4^poison^-mEos (Fig 4A).

**Figure 4.**
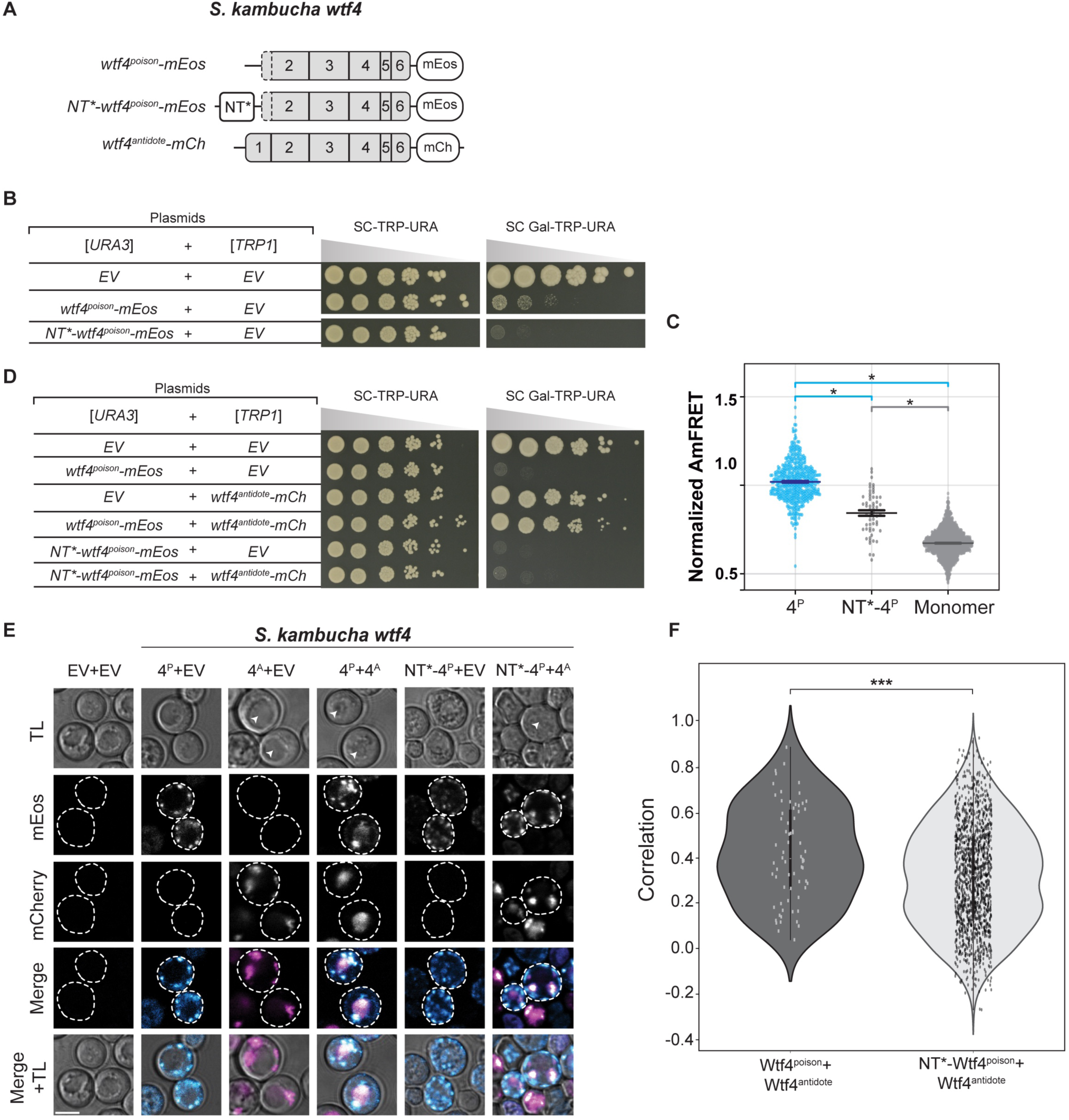
Reducing Wtf4^poison^ assembly with NT* tag increases toxicity and affects its rescue by Wtf4^antidote^. **A.** Cartoon of alleles used in this experiment (B-F). The NT* tag has a general anti-aggregation property [34]. **B.** A spot assay of cells serially diluted and plated on SC-TRP-URA and SC Gal- TRP-URA plates and grown at 30^·^C for 4 days. Each strain carries both a [*URA3*] and a [*TRP1*] plasmid. The plasmids are either empty (EV) or carry the indicated *wtf4* alleles under the control of galactose-inducible promoters. The horizontal break in the image for each plate is due to a rearrangement of the image to facilitate easy comparison. All strains were grown on the same plates (i.e., one SC-TRP-URA or SC Gal-TRP-URA plate). **C.** AmFRET values for three technical replicates of the specified Wtf4^poison^-mEos alleles and monomer-mEos (negative control). The median is indicated with a solid line and the bars represent the interquartile range. For easier comparison, the values were normalized so that Wtf4^poison^ had a median of 1 in each experiment. Since Wtf4-20^poison^-mEos cells were very low in number, we pooled two biological replicates (n=6 technical replicates). The data shown here do not include outliers. See S2 Data for the complete dataset and p-values. Statistical significance: *p<0.025, t-tests with Bonferroni correction. **D.** A spot assay of cells serially diluted and plated on SC-TRP-URA and SC Gal- TRP-URA plates and grown at 30^·^C for 3 days. Each strain carries both a [*URA3*] and a [*TRP1*] plasmid. The plasmids are either empty (EV) or carry the indicated *wtf4* alleles under the control of galactose-inducible promoters. All strains were grown on the same plates (i.e., one SC-TRP- URA or SC Gal-TRP-URA plate). **E.** Representative images of the same strains depicted in D were induced with galactose media for 4 hours at 30^·^C to express the indicated Wtf4^poison^-mEos proteins and/or Wtf4^antidote^-mCherry. The images are not at the same brightness and contrast settings to clearly show localization of tagged proteins. 4^P^ indicates Wtf4^poison^, 4^A^ indicates Wtf4^antidote^, TL is transmitted light, and the scale bar is 4 µm. **F**. Pearson’s Correlation between mCherry and mEos signal in cells from E expressing the specified proteins. N>100, *** p<0.001, t-test.

We found that this protein had a novel combination of assembly and localization not represented in any of our other alleles. Specifically, the NT*-Wtf4^poison^-mEos protein showed less self-assembly than Wtf4^poison^-mEos (Fig 4C), but the NT* tag did not visibly alter the localization of the protein within cells (Fig 4E). Interestingly, the NT*-Wtf4^poison^-mEos protein showed increased toxicity in cells (Fig 4B). To test if the effects of the NT* tag were specific to Wtf4^poison^ or more general, we also tagged *S. octosporus* Wtf25^poison^ with NT* to generate NT*- Wtf25^poison^-mEos (S6A Fig). As with NT*-Wtf4^poison^-mEos, we observed unaltered localization and increased toxicity of the NT*-Wtf25^poison^-mEos protein, relative to wild-type Wtf25^poison^-mEos (S6B and S6D Fig). We could not, however, quantify NT*-Wtf25^poison^-mEos assembly by DAmFRET as the high toxicity prevented us from obtaining sufficient viable cells with mEos fluorescence (S4E Fig).

We also assayed how another solubility tag, the *E. coli* Maltose Binding Protein (MBP), affected Wtf^poison^ toxicity [40–42]. The MBP-tagged Wtf4^poison^ (MBP-Wtf4^poison^-mEos) had the same phenotype as the NT*-tagged Wtf4^poison^ protein: decreased assembly, increased toxicity, unaltered localization, relative to the wild-type protein (S7 Fig). The phenotype of the MBP-tagged Wtf25^poison^ (MBP-Wtf25^poison^-mEos) however, did not mirror that of the NT*-tagged protein. Instead, we found that the MBP-Wtf25^poison^-mEos protein showed reduced, but still high toxicity. The MBP-Wtf25^poison^-mEos protein also showed an altered, stronger endoplasmic reticulum-like localization, as compared to the wild-type protein (S7A-C Fig). This protein thus adds to those described above in which we see more ER-like localization associated with reduced Wtf^poison^ toxicity. The MBP-Wtf25^poison^-mEos protein did exhibit self-assembly, but we could not compare it the wild-type protein as high toxicity limited the viable cells we could assay via DAmFRET (S4F and S7D Figs).

The bulk of our experiments are consistent with a model where the assembly properties and the distribution of Wtf^poison^ proteins within cells affects toxicity, with distributed, punctate assemblies exhibiting more toxicity. Increasing Wtf^poison^ solubility (while maintaining assembly), increases toxicity, unless localization is disrupted. Forcing Wtf^poison^ proteins into large, localized assemblies, suppresses toxicity. The non-toxic Wtf4Δ^10-poison^ protein that forms distributed assemblies, however, undermines this simple model or at least indicates that there are additional unidentified features that are essential for toxicity.

An additional possibility that we considered was that the expression levels of the Wtf^poison^ proteins likely contributes to the phenotypes we observed, despite expressing all alleles from a common vector backbone using a common promoter. To assay expression levels, we quantified fluorescence of the mEos tags shared by all alleles. We did not use western blots as that approach is challenging to apply to Wtf^poison^ proteins as they are toxic and highly hydrophobic [15,23]. Instead, we looked at the acceptor (red form of mEos) fluorescence in live cells analyzed in the DAmFRET experiments (S4 Fig; Data from Figs 2B, 4C, S2E and S3D). As expected, based on other studies [43–46], we observed significant fluorescent protein signal heterogeneity within cells expressing the monomer mEos control and each Wtf^poison^ protein tested (S4 Fig). We did not, however, find increased fluorescent signal in cells expressing the most toxic proteins (as determined by growth assays). On the contrary, we tended to observe less signal in the cells expressing the most toxic proteins (e.g., Wtf4^poison^ and NT*-Wtf4^poison^).

These data support a model in which cells expressing the higher levels of toxic proteins are more likely to be removed by death and are not quantified. As mentioned above, however, we also observed low signal in the cells expressing the non-toxic Wtf4-ex2Δ^poison^, Wtf4-TMD2Δ^poison^, and Wtf4-TMD6Δ^poison^ alleles (S4 Fig). The low levels of Wtf4-TMD6Δ^poison^ could be due to its degradation in the vacuole (S2F Fig). For the other two alleles, it is formally possible the low levels of these proteins in cells could contribute to their lack of toxicity.

### Limited modularity of the Wtf^antidote^-specific domain

We next wanted to test which protein features affect Wtf4^antidote^ function (Fig 1A and 1B). As described above, the Wtf4^antidote^ shares the resides encoded by exons 2-4 with the Wtf4^poison^ but has an additional N-terminal domain encoded by exon 1. We hypothesized that the amino acids encoded by exon 1 would be insufficient for function and that exons 2-6 would be required for protein self-assembly as they comprise the Wtf4^poison^ protein, which self-assembles [15]. As expected, we found that a protein consisting of only the exon 1-encoded residues linked to a mEos tag (Wtf4 Exon1-mEos) could not self-assemble (S8A and S8B Fig). The Wtf4 Exon1- mEos protein was also not trafficked to the vacuole, despite this protein harboring two PY motifs that promote ubiquitin-mediated trafficking of wild-type Wtf^antidote^ proteins [24] (S8E Fig). Finally, the Wtf4 Exon1-mEos protein could not rescue Wtf4^poison^ toxicity, which parallels recent results using distinct Wtf proteins (S8C and S8F Fig) [24]. These results demonstrate that the antidote-specific domain encoded by *wtf4* exon 1 is insufficient for antidote function.

To test if the exon 1 encoded domain was modular, we generated two mutants *wtf4^poison^-ex1* and *wtf4^poison^-ex1^int^*. The *wtf4^poison^-ex1* allele moves the antidote-specific domain moved to the C-terminus of the protein (S8A Fig). The *wtf4^poison^-ex1^int^*allele moved exon 1 a more central region of the protein (beginning of exon 4). This location is between the last two predicted transmembrane domains and is not predicted to disrupt them (Fig 1B). We found that mEos tagged versions of both Wtf4^poison^-ex1 and Wtf4^poison^-ex1^int^ proteins were both non-toxic and trafficked to the vacuole, suggesting they retained at least some antidote functionality (S8D and S8F Fig). Only the Wtf4^poison^-ex1, however, could neutralize the toxicity of Wtf4^poison^-mCherry (S8D and S8F Fig). These results demonstrate that the domain encoded by exon 1 is at least partially modular, but that a central location within the polypeptide can disrupt the Wtf^antidote^ protein’s ability to neutralize a Wtf^poison^. We speculate this could be because extensive, continuous, amino acid identity facilitates Wtf^poison^ and Wtf^antidote^ co-assembly.

#### Wtf^antidote^ requires more than physical linkage to effectively traffic Wtf^poison^ to the vacuole

We next wanted to determine if co-assembly serves merely to link Wtf^poison^ and Wtf^antidote^ proteins to allow for co-trafficking, or if co-assembly serves a more nuanced role in neutralizing Wtf^poison^ toxicity. To test this, we generated Wtf^poison^ and Wtf^antidote^ proteins that were physically linked, but not co-assembled, by artificially tethering Wtf^poison^ proteins to diverged Wtf^antidote^ proteins, with which they cannot co-assemble, using a combination of GFP and GFP-binding protein (GBP) tags (Figs 5A, 5B, S9A and S9B).

**Figure 5.**
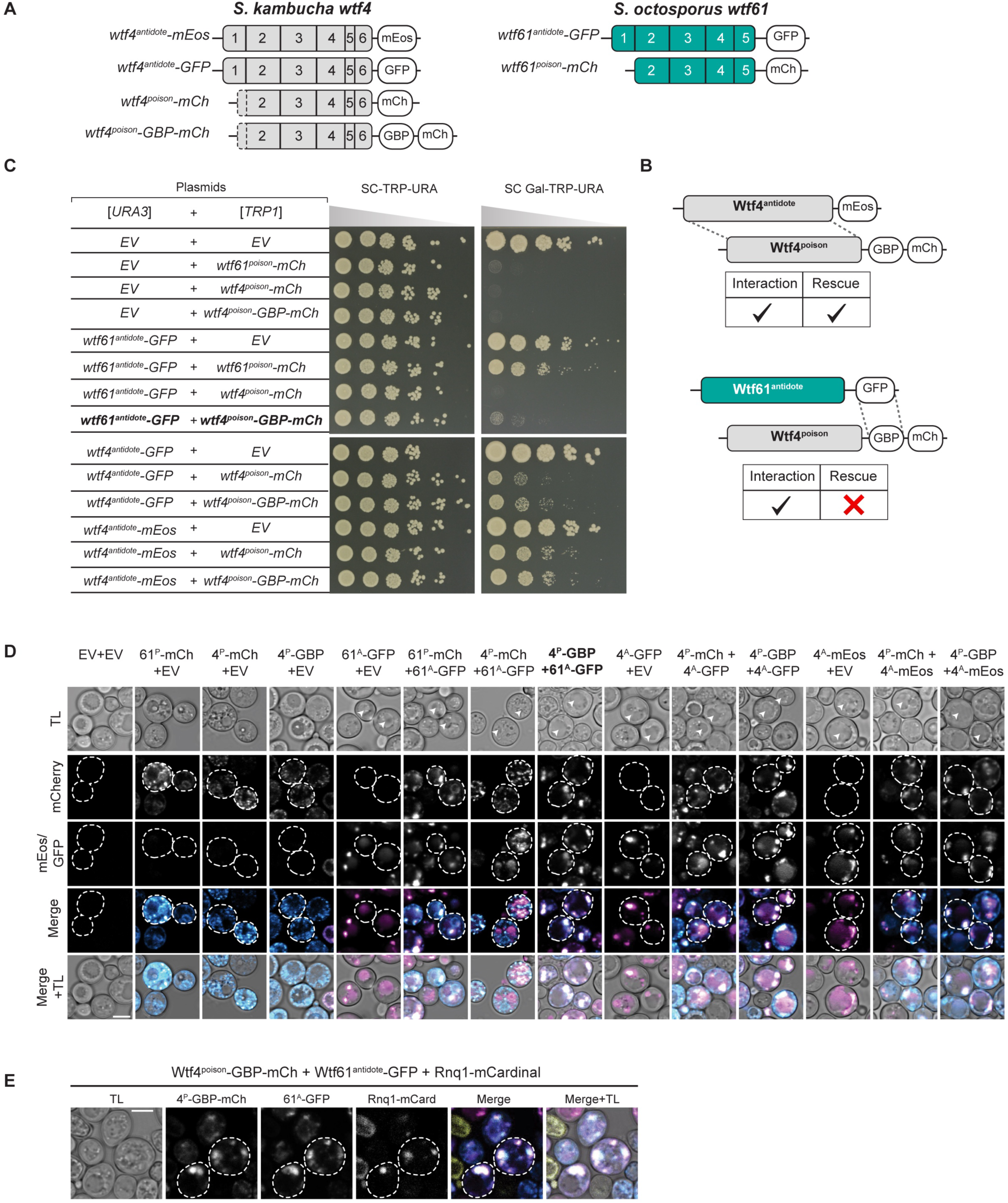
Effective neutralization of Wtf4^poison^ requires more than a physical connection to a Wtf^antidote^_._ **A.** Cartoon of constructs used in this experiment. *S. kambucha wtf4^poison^* was tagged at the C- terminus with either mCherry or GBP-mCherry (GBP: GFP-binding protein). *S. kambucha wtf4^antidote^*was tagged with mEos or GFP. *S. octosporus wtf61^antidote^* was tagged with GFP. *S. octosporus wtf61^poison^* was tagged with mCherry. **B.** Experimental set up and summary of the results shown in C and D. In a matching Wtf protein pair (top), poison-antidote interaction and rescue of poison toxicity is observed. In the mismatched pair (bottom), interaction between GFP and GBP results in a forced interaction between the poison and the antidote (shown in D). This interaction is insufficient to rescue the mismatched poison (shown in C). **C.** Spot assay of cells serially diluted and plated on SC-TRP-URA and SC Gal-TRP-URA plates and grown at 30^·^C for 3 days. Each strain carries both a [*URA3*] and a [*TRP1*] plasmid. The plasmids are either empty (EV) or carry the indicated *wtf* alleles under the control of galactose-inducible promoters. The horizontal break in the image of each plate is due to rearrangements of the images to facilitate easy comparison. All strains within a panel were grown on the same plates (i.e., one SC-TRP-URA or SC Gal-TRP-URA plate). **D.** Representative images of the same strains shown in C were induced in galactose for 4 hours at 30^·^C to express the indicated Wtf proteins. **E.** Representative images of cells induced with galactose and 500nM β-estradiol for 4 hours at 30^·^C to produce the indicated proteins. Rnq1-mCardinal marks the insoluble protein deposit (IPOD) that is associated with the vacuole [46–48]. In D-E, the images are not at the same brightness and contrast settings to clearly show localization of tagged proteins. 4^P^ indicates Wtf4^poison^, 4^A^ indicates Wtf4^antidote^, 61^P^ indicates Wtf61^poison^, 61^A^ indicates Wtf61^antidote^, TL is transmitted light, and the scale bar is 4 µm.

We first tested if tethering Wtf4^poison^ to *S. octosporus* Wtf61^antidote^ could neutralize the toxic Wtf4^poison^. We generated Wtf4^poison^-GBP-mCherry and Wtf61^antidote^-GFP proteins and found the tags did not disrupt protein function. Specifically, the Wtf4^poison^-GBP-mCherry protein was toxic, and could be rescued by Wtf4^antidote^ proteins, including Wtf4^antidote^-GFP. Similarly, the Wtf61^antidote^-GFP protein was able to rescue Wtf61^poison^-mCherry (Fig 5C). However, Wtf61^antidote^- GFP was not able to efficiently traffic Wtf4^poison^-GBP-mCherry to the vacuole or neutralize its toxicity (Fig 5C and 5D). This failure to rescue is not a failure of the GFP-GBP interaction to link the proteins as we found that the Wtf61^antidote^-GFP and Wtf4^poison^-GBP-mCherry proteins largely co-localized, which they did not do in the absence of the GBP tag (Fig 5D). Moreover, the localization of the tethered Wtf61^antidote^-GFP-Wtf4^poison^-GBP-mCherry changed relative to the individual proteins: the Wtf61^antidote^-GFP no longer trafficked to the vacuole and Wtf4^poison^-GBP- mCherry was less distributed in cells. Effective GFP-GBP linkage is the most parsimonious explanation of these observations. To test if our results were generalizable, we also analyzed a widely diverged, independent pair of Wtf proteins (*S. octosporus* Wtf25^poison^ and *S. cryophilus* Wtf1^antidote^). Our results with this protein pair mirrored those described above suggesting that specific antidote-poison co-assembly generally promotes proper antidote function (S9 Fig).

Given that the tethered Wtf61^antidote^-GFP and Wtf4^poison^-GBP-mCherry were often adjacent to vacuoles and not totally distributed in cells, as occurs when trafficking of Wtf proteins is disrupted [15], we suspected that the vacuolar targeting was still occurring, but blocked at a late step in the process (e.g., vacuole entry). Consistent with this notion, the assemblies often co- localized with the Rnq-1mCardinal protein, which is a prion protein that marks the insoluble protein deposit (IPOD) [47–49] (Fig 5E). This result was surprising to us because with a different expression system (β-estradiol induction) a considerable amount of the Wtf4^poison^/Wtf4^antidote^ co- assemblies accumulate at the IPOD are not toxic to cells [15]. It is unclear why GFP/GBP mediated assemblies, but not Wtf/Wtf mediated co-assemblies, would be toxic at the IPOD, but we speculate about this in the discussion.

Beyond the GFP/GBP tethering experiments, we made additional observations suggesting that a specific nature of co-assembly with Wtf^poison^ is required for Wtf^antidote^ function. When characterizing the toxic NT*-Wtf4^poison^ protein (with the NT* domain that increases solubility), we found that this protein was not effectively rescued by wild-type Wtf4^antidote^-mCherry (Fig 4D).

This drastic reduction in rescue (relative to that observed with the wild-type Wtf4^poison^) is striking given that the colocalization of NT*-Wtf4^poison^-mEos with Wtf4^antidote^-mCherry appears only mildly reduced relative to that observed between wild-type Wtf4 proteins (Figure 4E and 4F). In addition, the localization of the NT*-Wtf4^poison^-mEos is altered in the presence of Wtf4^antidote^- mCherry (it becomes less distributed compared to wild type Wtf4 proteins; Fig 4E). These observations suggest the NT*-Wtf4^poison^-mEos and Wtf4^antidote^-mCherry proteins interact, but are not efficiently trafficked into the vacuole (Fig 4E). This is analogous to the GFP/GBP linked, but unassembled, Wtf^poison^ and Wtf^antidote^ pairs described above. The effect of the NT* tag on Wtf^poison^/Wtf^antidote^ compatibility was not, however, universal. We did not observe strong disruption of poison/antidote compatibility caused by the NT* tag on *S. octosporus* Wtf25^poison^ (NT*-Wtf25^poison^ allele, S6 Fig), indicating that factors affecting poison/antidote compatibility can be context dependent.

Together, our results demonstrate that a physical linkage is insufficient to ensure efficient neutralization of a Wtf^poison^ protein’s toxicity by a Wtf^antidote^. Instead, specific co-assembly of the proteins likely both links compatible Wtf^poison^ and Wtf^antidote^ proteins and facilitates their effective co-trafficking into the vacuole.

### C-terminal region supports Wtf4^antdiote^ function

As introduced above, Hu et al [17] previously described mutant alleles of two *S. pombe wtf* genes lacking the codons for the last 10 amino acids. Those mutants maintained Wtf^antidote^ activity [17]. To test if the C-terminal amino acids were generally dispensable for antidote function, we made *wtf4^antidote^*truncation alleles. We found that a 10 amino acid truncation of Wtf4, (Wtf4Δ^10-antidote^-mCherry) was slightly toxic to cells, as compared to the Wtf4^antidote^-mCherry and empty vector controls (S10 Fig). A 20 amino acid truncation of the, Wtf4Δ^20-antidote^-mCherry, showed even more toxicity, although it was still considerably less than the toxicity observed with the Wtf4^poison^-mCherry. A 29 amino acid truncation, Wtf4-ex6Δ^antidote^-mCherry, exhibited no toxicity (S10 Fig). These results suggest that the C-terminal 20 amino acids of Wtf4^antidote^ play a role in limiting the toxicity of the Wtf4^antidote^ protein.

We next tested if the truncation Wtf4^antidote^ proteins could rescue the toxicity of a wild-type Wtf4^poison^-mEos protein. Neither the Wtf4Δ^10-antidote^-mCherry or Wtf4-ex6Δ^antidote^-mCherry proteins appreciably neutralized Wtf4^poison^-mEos toxicity (S10A and S10B Fig). The Wtf4Δ^20-antidote^- mCherry protein, however, rescued growth of cells expressing Wtf4^poison^-mEos to a level comparable to that observed in the cells expressing only Wtf4Δ^20-antidote^-mCherry. So, despite the slight toxicity of the Wtf4Δ^20-antidote^-mCherry protein, it still retained some antidote function.

Interestingly, we were surprised to see that the other truncated antidotes Wtf4Δ^10-antidote^-mCherry and Wtf4-ex6Δ^antidote^-mCherry could also partially rescue the toxicity of Wtf4Δ^20-poison^-mEos poison, although it was much less rescue compared to that of the wild-type Wtf4^antidote^-mCherry protein (S10A and S10B Fig). This demonstrates that these truncated proteins retain some functionality.

Our results indicate that the C-terminus supports Wtf4^antidote^ function, but that some functionality remains even in the absence of the last 29 amino acids. When considered in combination with the results of Hu et al [17], our combined results indicate that the importance of the C-terminus is context dependent. Like our Wtf^poison^ results discussed earlier, these observations, combined with the lack of conservation of Wtf proteins, suggests that Wtf^antidote^ function (outside of the conserved PY motifs) relies on overall properties of the protein (e.g., lack of toxicity and ability to co-assemble with a matching Wtf^poison^), rather than any functional domain.

#### Self-killing alleles that encode a functional Wtf^poison^ with an incompatible Wtf^antidote^

Because Wtf proteins are encoded on largely overlapping coding sequences, a change in the shared coding sequence can simultaneously create a novel Wtf^poison^ protein and a matching novel Wtf^antidote^ protein. Our engineered mutant alleles offered the opportunity to explore if matching proteins are always compatible (i.e., if a novel toxic Wtf^poison^ protein is always neutralized by its corresponding Wtf^antidote^ protein).

We therefore explored poison and antidote compatibility more broadly within the alleles we generated. In many cases, the Wtf^antidote^ proteins were able to rescue their matching Wtf^poison^ alleles (S1 Table). One class of mutants, however, created toxic Wtf^poison^ proteins that were not rescued by the matching Wtf^antidote^. In *wtf4*, these mutants changed a region of exon 6 that encodes a 7 amino acid repeat found in variable numbers (0-84 base pairs of repeat sequence) within *S. pombe wtf* genes (Fig 6A and 6B) [20]. The repeat region in *wtf4* encodes one complete repeat plus three additional amino acids of the repeat (Fig 6C). We found that mutating this region in Wtf4 by either scrambling the amino acid order (Wtf4-rep2^sc^) or by replacing the ten amino acids with alanine (Wtf4-rep2^A^ allele) generated toxic Wtf^poison^ proteins that were not neutralized by their matching Wtf^antidote^ proteins (Fig 6D). The Wtf4-rep2^sc-antidote^- mCherry and Wtf4-rep2^A-antidote^-mCherry proteins were both trafficked to the vacuole when expressed alone but did not effectively co-traffic their corresponding Wtf^poison^ proteins (Fig 6E).

**Figure 6.**
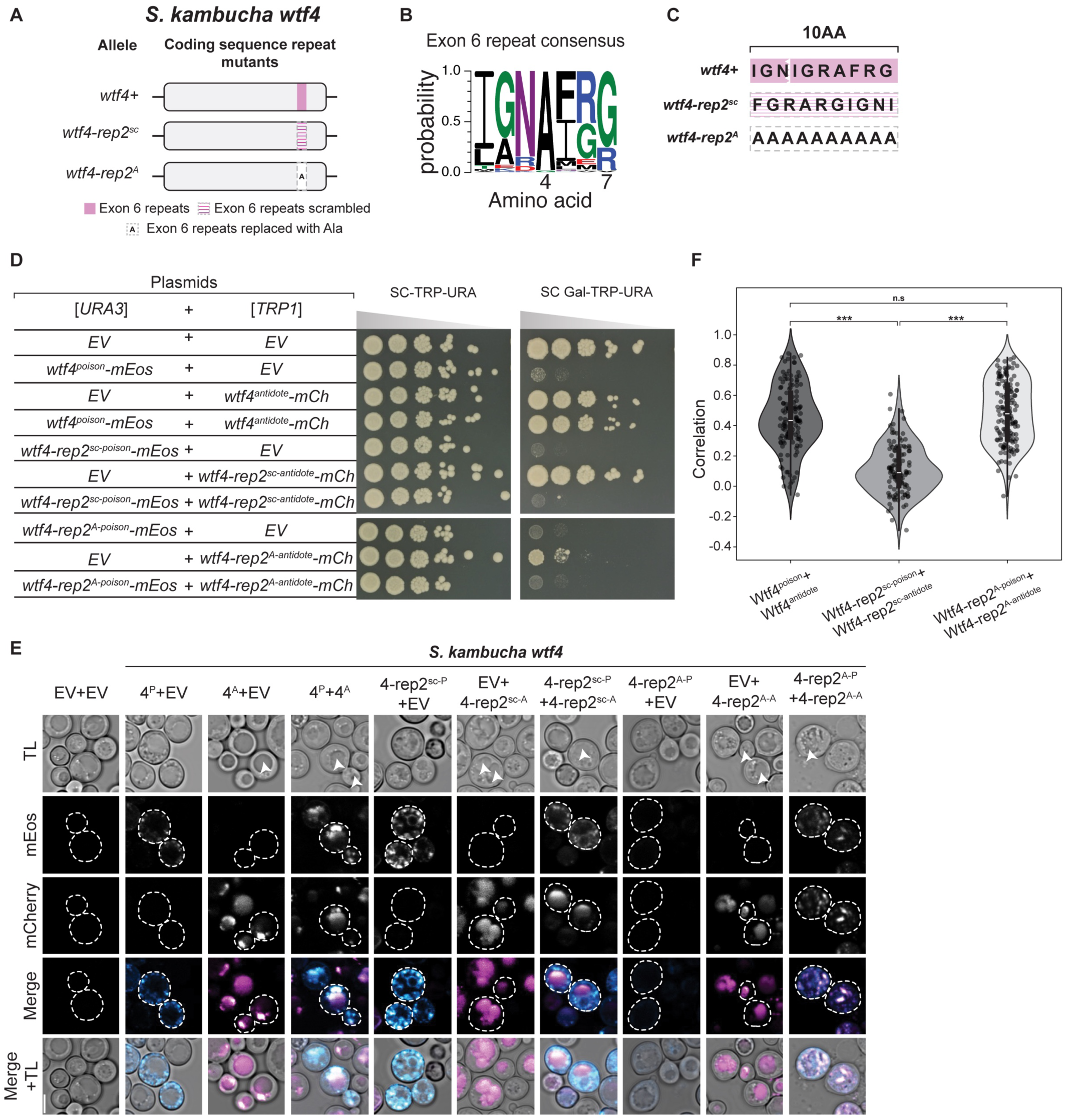
Modification of *wtf4* exon 6 CDS repeats can disrupt antidote rescue. **A.** Cartoon of two coding sequence repeat mutants of *S. kambucha wtf4*. **B**. Logo representing the amino acids encoded by the repeats found in exon 6 of *S. pombe wtf* genes from [20]. **C.** The amino acids encoded by the exon 6 repeats in *S. kambucha wtf4* and the *wtf4-rep2^sc^*and *wtf4-rep2^A^* mutant alleles. **D.** Spot assay of cells serially diluted and plated on SC-TRP-URA and SC Gal-TRP-URA plates and grown at 30^·^C for 3 days. Each strain carries both a [*URA3*] and a [*TRP1*] plasmid. The plasmids are either empty (EV) or carry the indicated *wtf4* alleles under the control of galactose-inducible promoters. The horizontal breaks in the image of each plate are due to rearrangements of the images to facilitate easy comparison. All strains within a panel were grown on the same plates (i.e., one SC-TRP-URA or SC Gal-TRP-URA plate). **E.** Representative images the same strains depicted in D were induced in galactose for 4 hours at 30^·^C to express the indicated Wtf proteins. The images are not at the same brightness and contrast settings to clearly show localization of tagged proteins. The arrows in the TL panels highlight vacuoles. 4^P^ indicates Wtf4^poison^, 4^A^ indicates Wtf4^antidote^, TL indicates transmitted light, and the scale bar is 4 µm. **F**. Pearson’s Correlation between mEos and mCherry signal in cells expressing the specified constructs from E. N>100, ***p<0.001, t-test.

**Figure 7.**
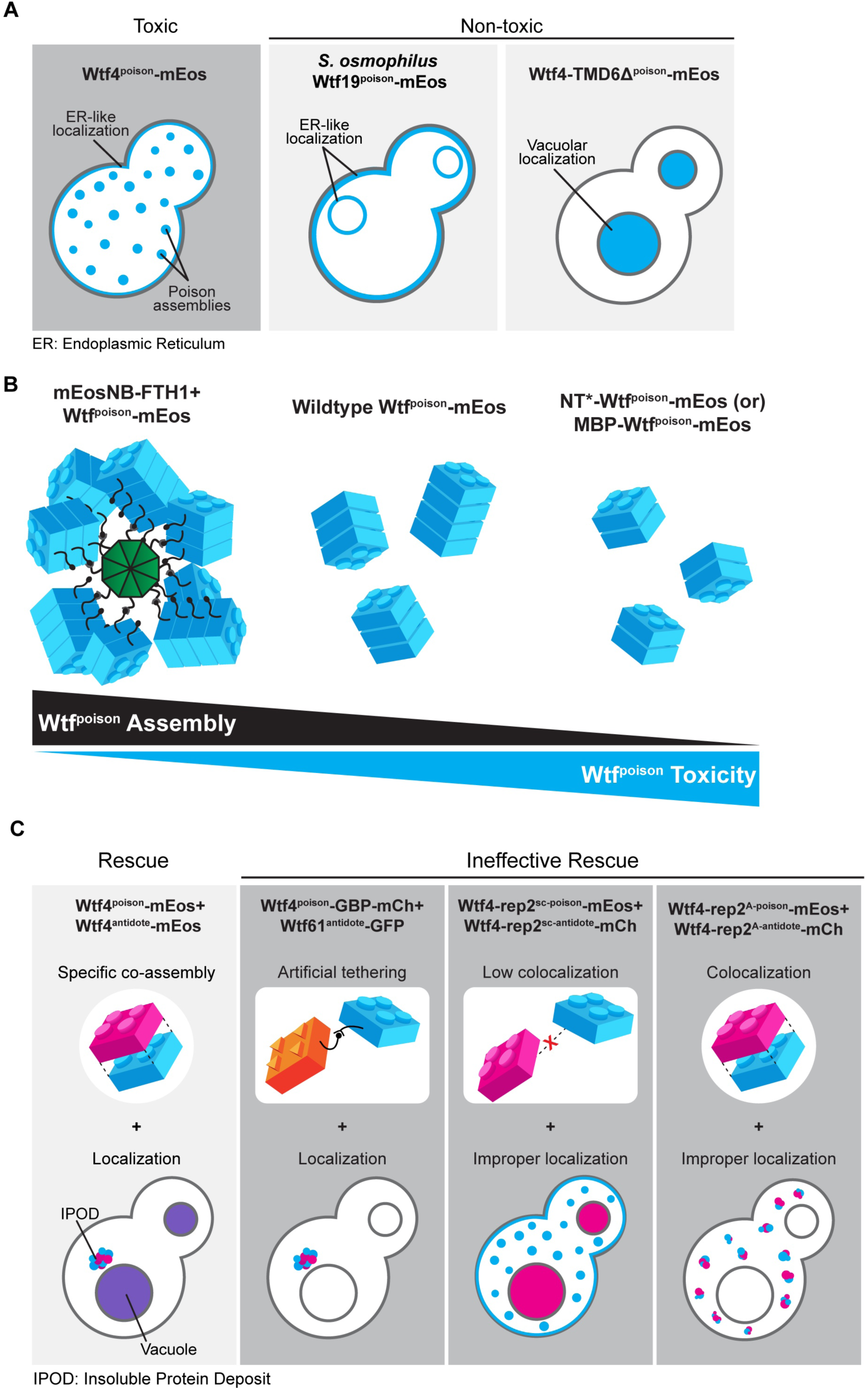
Analyses of Wtf proteins reveal additional functional constraints. **A.** A dispersed Wtf^poison^ localization and poison-poison assembly can be observed across greatly diverged proteins and are associated with Wtf^poison^ toxicity. However, assembly and/or localization alone are insufficient to cause toxicity. In multiple examples of non-toxic proteins (an example is included in each panel), we observe either an endoplasmic reticulum-like localization or vacuolar localization. **B.** Modulating Wtf^poison^ assembly with exogenous tags affects its toxicity. While increasing Wtf^poison^ assembly via tethering to mEosNB-FTH1 suppresses toxicity, increasing solubility with NT* (or MBP in the case of *S. kambucha* Wtf4^poison^) increases the toxicity. **C.** Multiple modalities of ineffective antidote rescue. Effective rescue of a Wtf^poison^ by the corresponding Wtf^antidote^ (e.g., Wtf4 proteins) is supported by specific co-assembly and vacuolar localization. Artificial tethering of a mismatched Wtf^poison^ and Wtf^antidote^ does not result in effective rescue. Disruption of poison-antidote co-localization (e.g., Wtf4-rep2^sc^ proteins) or vacuolar targeting of co-localized poison-antidote proteins (e.g., Wtf4-rep2^A^ proteins) also results in ineffective rescue.

The Wtf4-rep2^sc^ ^antidote^-mCherry and Wtf4-rep2^sc^ ^poison^-mEos proteins showed decreased co- localization, relative to wild-type proteins, suggesting the proteins had disrupted co-assembly (Fig 6E and 6F). Alternatively, the Wtf4-rep2^A^ ^antidote^-mCherry and Wtf4-rep2^A^ ^poison^-mEos proteins colocalized (Fig 6F) but remained more distributed within cells relative to wild-type (Fig 6E). This distributed localization is a change from the vacuolar localization of Wtf4-rep2^A^ ^antidote^-mCherry alone, further supporting that the Wtf4-rep2^A^ ^antidote^-mCherry and Wtf4-rep2^A^ ^poison^-mEos are interacting. Therefore, the incompatibility of the Wtf4-rep2^A^ proteins adds additional support to our earlier conclusion that a particular form of association is required between Wtf^antidote^ and Wtf^poison^ proteins to ensure the poison is effectively trafficked to the vacuole and neutralized.

We also assayed the effects of deleting the repeat region of *wtf4* exon 6 alone (*wtf4-rep2Δ* allele), or in combination with deleting another repetitive region found in exon 3 (*wtf4-rep1-2Δ* allele; S11A Fig) [20]. We found that the Wtf4-rep2Δ^poison^-mEos protein is toxic but is only partially rescued by Wtf4-rep2Δ^antidote^-mCherry, relative to the rescue observed between wild-type Wtf4 poison and antidote proteins (S11D Fig). We found that the two Wtf4-rep2Δ proteins exhibit decreased colocalization, relative to wild-type proteins, suggesting the limited rescue is due to disrupted poison-antidote interaction (S11E and S11F Fig). Interestingly, we found that the defect in poison-antidote compatibility conferred by the deletion of the exon 6 repeats in the *wtf4-rep2Δ* allele is partially suppressed by also deleting the repetitive region found in exon 3. Specifically, the proteins encoded by the *wtf4-rep1-2Δ* allele, with both regions deleted, have near wild-type phenotypes (S11 Fig). We observed no defects in poison and antidote compatibility in the proteins encoded by an allele (*wtf4*-*rep1Δ*) lacking only the repetitive region in exon 3 (S11A, S11B, S11D-F Fig).

Finally, we also tested if the novel mutations we made in this study in the exon 3 or C-terminal repeats affected compatibility with wild-type Wtf proteins. In all cases tested, the repeat mutant Wtf^poison^ proteins were not neutralized by their wild-type Wtf^antidote^ counterparts and vice versa (S12 Fig). For example, Wtf4-rep1Δ^poison^-mEos is not neutralized by Wtf4^antidote^-mCherry and Wtf4^poison^-mEos is not neutralized by Wtf4-rep1Δ^antidote^-mCherry (S12B Fig).

The repeats in exon 6, but not in exon 3, are broadly conserved in the *wtf* gene family [19]. We also mutated the homologous region in *S. octosporus wtf25* and found analogous phenotypes as those described above in *wtf4* (S13 Fig). Like *wtf4*, *S. octosporus wtf25* also encodes 10 amino acids in this C-terminal region (S13A-C Fig). We generated the *S. octosporus wtf25-rep2^Sk^* allele by swapping the endogenous codons for those of *wtf4*. We found that this allele encoded an incompatible Wtf25-rep2*^Sk^*^-poison^-mCherry and Wtf25-rep2*^Sk^*^-antidote^-mEos pair (S13D Fig). Surprisingly, the individual Wtf25-rep2*^Sk^* proteins were compatible with Wtf25 proteins (i.e., Wtf25-rep2*^Sk^*^-poison^-mCherry was rescued by Wtf25^antidote^-mEos and vice versa; S13D Fig). We also deleted the repeat region of *S. octosporus wtf25* (to generate the *wtf25-rep2Δ* allele) and again found phenotypes similar to those observed in the analogous *wtf4* mutant (*wtf4-rep2Δ*) (S14 Fig). Specifically, the Wtf25-1^antidote^-mCherry showed reduced rescue of the Wtf25- rep2!1^poison^-mEos toxicity, relative to the wild-type Wtf25 protein pair (S14C Fig).

Altogether, our results further support a critical role for the repeats in Wtf^poison^ and Wtf^antidote^ compatibility and reveal that mutants in this domain can encode Wtf^poison^ proteins not neutralized by the matching Wtf^antidote^ proteins. Such alleles are important constraints on *wtf* gene evolution as they would contribute to infertility via self-killing.

## Discussion

### Protein assembly plays conserved roles in Wtf protein function

The mutant analyses provided in this work expands and supports a working model in which 1) Wtf^poison^ toxicity is tied to the homotypic assembly of the proteins [15], 2) Wtf^antidote^ proteins are ubiquitinated and trafficked to the vacuole [15,24] and, 3) Wtf^antidote^ proteins co-assemble with their matching Wtf^poison^ proteins and co-traffic them to the vacuole [15,26]. Recent work has established that ubiquitin-mediated Wtf^antidote^ trafficking and co-trafficking with their corresponding Wtf^poison^ proteins are conserved within the extremely diverse gene family [24]. It was unclear, however, if homotypic protein assembly is a conserved feature of Wtf^poison^ or Wtf^antidote^ protein function.

The results of this study support the model that homotypic protein assembly plays critical roles in the function of Wtf^poison^ proteins. First, we observed that the ability to self-assemble was conserved amongst functional (i.e., toxic) wild-type Wtf^poison^ proteins from four *Schizosaccharomyces* species (Fig 2). Given that the functional proteins share as little as 20% amino acid identity, these observations likely reflect conserved functional importance of self-assembly. Our mutant analyses of Wtf4^poison^ proteins provided additional support for this model in that all toxic mutant proteins self-assembled (S2 and S3 Figs). Furthermore, we found that the Wtf^poison^ toxicity could be modulated by altering the assembly properties of the protein with tags (Figs 3, 4, S6 and S7). Together, our data suggest that Wtf^poison^ toxicity is tied to protein assembly with distributed, small assemblies showing greater toxicity than localized, larger assemblies (Figs 2-4, 7, S2, S3, S6 and S7). The recent work of Zheng et al. [24] suggests the distributed assemblies may represent localization to the trans-Golgi network.

Still, it is important to note that the Wtf4Δ^10^ ^poison^ allele was non-toxic, despite assembling into small, distributed assemblies indistinguishable from those generated by the wild-type protein (S3 Fig). This exceptional protein highlights that even if self-assembly is critical for Wtf^poison^ toxicity, it is not the only factor required. In addition, this allele offers an opportunity for future work to explore features that distinguish toxic from nontoxic protein assemblies.

Our results also support an expanded, more nuanced role for homotypic protein assembly and trafficking in Wtf^antidote^ function. First, we found that the antidote-specific domain that contains the PY motifs is insufficient to promote vacuole trafficking, suggesting some other features of Wtf proteins are also required (S8 Fig). We posit protein self-assembly could contribute to this function. For Wtf^antidote^ neutralization of a Wtf^poison^, we initially assumed that co-assembly of Wtf^antidote^ with Wtf^poison^ proteins served only to physically link the proteins, thus enabling the Wtf^antdiote^ to traffic the Wtf^poison^ to the vacuole [15]. We found, however, that physical linkage between a Wtf^poison^ and Wtf^antidote^ can be insufficient to ensure their co-trafficking to the vacuole and neutralization of the Wtf^poison^’s toxicity. We observed this insufficiency in experiments linking two distinct pairs of non-matching Wtf^poison^ and Wtf^antidote^ proteins with GFP-GBP tags (Figs 5 and S9). In addition, we found that the *wtf4-rep2^A^* and *NT*-wtf4* alleles encode protein pairs in which poison-antidote interaction appeared largely intact, but trafficking into the vacuole and poison neutralization was disrupted (Figs 4 and 6). These experiments suggest that efficient co- trafficking of Wtf^poison^/ Wtf^antidote^ protein assemblies requires a particular conformation or strong affinity between the interacting proteins not replicated in the ineffective Wtf^poison^/Wtf^antidote^ combinations mentioned above (Fig 7).

Moreover, our experiments surprisingly revealed that trafficking toxic Wtf assemblies to the proper destination may not always be sufficient to ensure their neutralization. In previous work using a different induction system than that employed here, we found that much of the trafficked Wtf4^poison^/Wtf4^antidote^ assemblies accumulated at the insoluble protein deposit (IPOD), in addition to the vacuole [15] and the cells were viable. Similar, but generally smaller, assemblies can be seen accumulating outside of the vacuole (likely the IPOD) in many of the viable cells we image expressing Wtf^antidote^ proteins or compatible Wtf^poison^/Wtf^antidote^ assemblies using the GAL induction system employed in this study (e.g., Fig 6E panels expressing Wtf4^antidote^ and Wtf4^antidote^/Wtf4^poison^). Similarly, Zheng et al [24] found that vacuole localization was not essential for Wtf^poison^ neutralization as they observed that trafficking to the endosome can be sufficient.

Here, however, we observed that Wtf4^poison^-GBP/Wtf61^antidote^-GFP assemblies (and likely the Wtf25^poison^-GBP/Wtf1^antidote^-GFP assemblies) were trafficked to the IPOD, but were still toxic (Figs 5, 7 and S9). These results suggest that factors beyond localization, perhaps assembly conformation, can also affect the toxicity of Wtf proteins.

### Commonalities between Wtf proteins and other self-assembling proteins

The Wtf proteins share broad parallels with other nonhomologous proteins that form assemblies. For example, a sequence-independent common oligomeric property may underly the toxicity of unrelated amyloid proteins [50]. While Wtf^poison^ proteins are related to each other, their sequences are extremely diverged. We propose a common feature of their assembled forms is likely responsible for their shared toxicity. Another feature Wtf proteins share with several unrelated proteins is the capacity to form functional assemblies. Multiple amyloidogenic proteins form functional amyloids that perform diverse biological functions, including long term memory in flies [51], epigenetic inheritance in yeast [52] and biofilm formation in bacteria [53–55]. Some functionally aggregating amyloids have also shown to be toxic at intermediate stages of assembly, suggesting that protein toxicity could pose a risk in certain instances [56]. The co- expression of Wtf^antidote^ with the toxic Wtf^poison^ results in a change in its localization and the assembly properties of the Wtf^poison^-Wtf^antidote^ complex. This suggests that protein-protein interaction within a pair of corresponding Wtf proteins is similar to functional aggregation. While the outcome of functional Wtf protein assembly is different (i.e., successful drive), the delicate interplay between toxic and non-toxic protein assemblies is similar to functional aggregating amyloids.

The use of E3 ubiquitin ligases to direct protein trafficking is an additional theme Wtf proteins share with other self-assembling proteins acting in diverse cell signaling processes, including immune response [57,58], prion disease [59] and other neurodegenerative disorders [60–66]. In multiple cases, ubiquitination of a key protein results in its aggregation, differential trafficking to specific intracellular locations or degradation, suggesting that this is a common mechanism for enabling downstream signaling processes. Additionally, a lack of ubiquitination by the ligase often results in toxic aggregates [59,62,65] or reduced functionality of the key protein [57,58], which are very reminiscent of the Wtf^poison^/Wtf^antidote^ assemblies discussed above.

### Rapidly evolving coding sequence repeats can affect Wtf^poison^-Wtf^antidote^ compatibility

Most *S. pombe wtf* genes contain varying copy numbers of a sequence repeat in exon 3 [20]. The potential role of the exon 3 repeats was unclear, but all functionally validated *S. pombe wtf* drivers contain the exon 3 repeats (S2 Table). One *wtf* gene (*wtf23* from the CBS5557 strain) that lacks the exon 3 repeats has been tested in *S. pombe* and failed to cause drive in two strain backgrounds, although it was not determined if the encoded proteins were non-functional or if the driver was effectively suppressed [20,25]. Previous work also found that a mismatch of repeat numbers in exon 3 between a Wtf13^poison^ (five repeats) and Wtf18-2^antidote^ (four repeats) could still produce a compatible poison and antidote pair [26]. In this work, we found that deleting the exon 3 repeats in *wtf4* (*wtf4-rep1Δ* allele) produced a functional, compatible Wtf^poison^ and Wtf^antidote^ pair. The Wtf4-rep1Δ^poison^, however, was not neutralized by the wild-type Wtf4^antidote^, which has two repeats (S12B Fig). This indicates that the exon 3 repeats can affect Wtf^poison^ and Wtf^antidote^ compatibility in a context dependent fashion.

Many *S. pombe wtf* genes also contain a sequence repeat in exon 6 [20]. Repeats homologous to those found in *S. pombe* exon 6 can also be found in the C-termini of many genes in *S. octosporus* and *S. osmophilus* [19]. All functionally validated drivers contain repeats in *S. octosporus*, but one functional *S. pombe* driver (*wtf35* from FY29033) lacks the repeats (S5 Table) [19,25]. Previous work demonstrated that a mismatch in the number of exon 6 repeats could affect Wtf^poison^ and Wtf^antidote^ compatibility in *S. pombe* Wtf proteins [15,26]. This current work extends previous work by showing with additional alleles of *wtf4* and novel alleles of *S. octosporus wtf25* that copy number mismatches in this region disrupt Wtf^poison^ and Wtf^antidote^ compatibility.

Overall, our results support the model that the rapid copy number evolution of the repeats found in *wtf* genes contribute to rapid innovation of novel Wtf^poison^ and Wtf^antidote^ pairs [19,20]. These highly evolvable sequences have likely contributed to the long-term evolutionary success of *wtf* drivers, as generating novel alleles allows frequent generation of novel drivers likely to be heterozygous (and thus drive) in crosses. Frequent driver turnover may also complicate the evolution of drive suppressors that are not other *wtf* genes. Still, this work reveals that these hypermutable regions come with a burden as they contain the potential to generate self-killing alleles, which are discussed below.

### Wtf fitness landscape likely includes self-killing alleles

Our analyses of mutations in the repeats found in exon 6 of *S. pombe* genes revealed a novel self-killing phenotype in which a *wtf* allele can encode a toxic Wtf^poison^ that is not effectively neutralized by its corresponding Wtf^antidote^. Such an allele is expected to lead to a dominant loss of fertility, analogous to mutations where the antidote protein expression or function is disrupted [17,22,24]. This phenotype was strongest in mutants that changed the sequence of the repeats (i.e., *wtf4-rep2^sc^*, *wtf4-rep2^A^*and *wtf25-rep^Sk^*). These mutations are rather dramatic and have a low probability of arising spontaneously in nature.

A weaker version of the self-killing phenotype was, however, also observed in mutations that deleted the repeats. Interestingly, the deleterious effects of removing the exon 6 repeats in *wtf4* (*wtf4-rep2Δ* allele) could be suppressed by also deleting the exon 3 repeats (*wtf4-rep1-2Δ* allele) (S12C and S12D Fig). This suppression, in addition to the one functional driver known to lack the exon 6 repeats, shows that changes in other regions of the protein can compensate for the missing repeats. Still, the existence of *wtf* genes without the repeats, and extensive gene conversion within the family, suggests that novel deleterious repeat deletion mutations are likely to arise recurrently in natural populations [19–21].

We may have also fortuitously sampled one largely self-killing allele from a natural population that did not have disrupted repeats, suggesting there are multiple paths to generating such alleles. The *wtf41* gene from *S. osmophilus* encodes a toxic Wtf41^poison^ that is not efficiently neutralized by the corresponding Wtf41^antidote^ protein (S5C Fig, [19]). All together, we propose self-killing *wtf* alleles could contribute to recurrent, spontaneous sub-fertility or infertility. We propose this spontaneous sub-fertility could be a persistent burden on the population fitness of all *Schizosaccharomyces* species carrying *wtf* drivers.

### Rapid evolution and the risk of self-killing alleles

Beyond *wtf* genes, there are many known killer meiotic drivers [3,5,6,8,9,11]. There are also likely many more yet to be discovered, given their accelerated pace of discovery in recent years. Although the genes causing drive in different systems are not homologous, they often share mechanistic and evolutionary themes [13,14,67]. Those themes include production of poisons/killer elements and rapid evolution. The *wtf* genes have illustrated that high evolvability, via nonallelic gene conversion and mutable coding sequence repeats, can facilitate the evolutionary success of meiotic drivers [19,21,68]. This work reveals that rapid evolution of killer elements also presents the risk of generating self-killing alleles. There is no reason to suppose such risks would be specific to the *wtf* killers. Instead, we posit that such self-killing alleles may be a widespread source of recurrent, spontaneous infertility in eukaryotes.

## Materials and Methods

### Cloning

We confirmed all the vectors described in this study by Sanger sequencing or by Nanopore sequencing via Plasmidsaurus. The specifics for the yeast strains used in this study are listed in S4 Table, plasmids are in S5 Table and the oligos are in S6 Table.

#### S. cerevisiae vectors

##### Generation of a Gal-inducible GFP vector to tag alleles

We amplified GFP-ADH1 terminator from a previously published plasmid, pSZB464 [15] with oligos 3744+3743. We digested this product with SacI and SpeI and cloned it into pDK20 [69] to generate pSZB1528. We then isolated the Gal promoter-GFP-ADH1T after digestion with SacI and KpnI and cloned it into a SacI-KpnI digested pRS316 [70] to generate pSZB1540.

##### Generation of a Gal-inducible mCherry vector to tag alleles

We amplified mCherry-ADH1 terminator from pFA6a-mCherry-kanMX6 [71] with oligos 3745+3743. We digested this product with SacI and SpeI and cloned it into pDK20 [69] to generate pSZB1526. We then isolated the Gal promoter-mCherry-ADH1T after digestion with SacI and KpnI and cloned it into a SacI-KpnI digested pRS314 [70] to generate pSZB1537.

##### Generation of a Gal-inducible mEos3.1 vector to tag alleles

We digested V08 [30] with SacI and KpnI to release the fragment with Gal promoter- 4x(EAAAR) linker-mEos3.1. We then cloned this into SacI-KpnI digested pRS316 [70] to generate pSZB1460.

##### Generation of a monomer mEos3.1 in an ARS/CEN plasmid

We cut out Gal-monomer mEos3.1-cyc1T from RHX0935 [30] with SacI and KpnI, and cloned into SacI, KpnI cut pRS316 [70] to generate pSZB1514.

##### Generation of Gal-inducible wtf alleles

###### S. kambucha wtf4 exon 1

The 136 base pairs that makes up exon 1 of *S. kambucha wtf4* was synthesized and cloned into pSZB1460 by IDT to generate pSZB1552.

###### S. kambucha wtf4-rep1Δ

We deleted 66 base pairs (bases 313-378 of poison coding sequence) that make up the exon 3 coding sequence repeats in *S. kambucha wtf4^poison^* and cloned it into pSZB1460 to generate pSZB1565. We repeated the same deletion in *S. kambucha wtf4^antidote^*(bases 439-504 of antidote coding sequence) and this construct was synthesized and cloned it into pSZB1537 by IDT to generate pSZB1736.

###### S. kambucha wtf4-rep2Δ

We deleted 30 base pairs (bases 802-831 of poison coding sequence) that make up the exon 6 repeats in *S. kambucha wtf4* and this construct was synthesized and cloned it into pSZB1460 to generate pSZB1566 by IDT. To construct the mutant antidote, we deleted bases 928-957 of the antidote coding sequence and this construct was synthesized and cloned it into pSZB1537 by IDT generate pSZB1737.

###### S. kambucha wtf4-rep1-2Δ

We deleted 66 base pairs (bases 313-378 of poison coding sequence) and 30 base pairs (bases 802-831 of poison coding sequence) to delete both the exon 3 and 6 coding sequence repeats in *S. kambucha wtf4.* This construct was synthesized and cloned into pSZB1460 to generate pSZB1670. To construct the mutant antidote, we deleted 66 base pairs (bases 439-504 of antidote coding sequence) and 33 base pairs (bases 928-957 of antidote coding sequence). This construct was synthesized and cloned it into pSZB1537 by IDT generate pSZB1738.

###### S. kambucha wtf4-rep2^sc^

We randomly scrambled the 10 amino acids that make up the exon 6 coding sequence repeats in *S. kambucha wtf4^poison^* and then reordered the codons to match the amino acids. We then replaced the wild-type 30 base pairs with the scrambled 30 base pairs (TTTGGGAGAGCGAGAGGGATAGGTAATATA) and this construct was synthesized and cloned into pSZB1460 to generate pSZB1742. To construct the mutant antidote, we replaced the exon 6 coding sequence repeats with the same scrambled 30 base pairs, and this construct was synthesized and cloned into pSZB1537 by IDT to generate pSZB1740.

###### S. kambucha wtf4-rep2^A^

We replaced the 30 base pairs that make up the exon 6 coding sequence repeats with alanine codons (GCAGCGGCTGCCGCTGCAGCTGCCGCAGCG) in *S. kambucha wtf4^poison^* and this construct was synthesized and was cloned into pSZB1460 to generate pSZB1743. To construct the mutant antidote, we replaced the exon 6 coding sequence repeats with the same alanine codons, and this construct was synthesized and cloned into pSZB1537 by IDT to generate pSZB1741.

###### S. kambucha wtf4^poison^-ex1^int^

We inserted exon 1 in between exons 3 and 4 of *S. kambucha wtf4^poison^* and this construct was synthesized and cloned into pSZB1460 by IDT to generate pSZB1616. To maintain the in-frame codons, we inserted exon 1 at 541 base pairs of the poison coding sequence, which is one base pair before exon 3 ends.

###### S. kambucha wtf4^poison^-ex1

We inserted exon 1 before the stop codon of *S. kambucha wtf4^poison^* coding sequence and this construct was synthesized and cloned into pSZB1460 by IDT to generate pSZB1555.

###### S. kambucha wtf4-TMD1Δ

We used TMHMM2.0 [71,72] to predict transmembrane topology (see S3 Table for a detailed description). With these predictions as guidance, we deleted the first predicted transmembrane domain (bases 121-186 of poison coding sequence) from in *S. kambucha wtf4^poison^* and this construct was synthesized and cloned into pSZB1460 by IDT to generate pSZB1561.

###### S. kambucha wtf4-TMD2Δ

We deleted the second predicted transmembrane domain (bases 232-291 of poison coding sequence) of *S. kambucha wtf4^poison^* coding sequence and this construct was synthesized and cloned into pSZB1460 by IDT to generate pSZB1562.

###### S. kambucha wtf4-TMD6Δ

We deleted the sixth predicted transmembrane domain (bases 580- 648 of poison coding sequence) of *S. kambucha wtf4^poison^* coding sequence and this construct was synthesized and cloned into pSZB1460 by IDT to generate pSZB1563.

###### S. kambucha wtf4-ex2Δ

We deleted exon 2 (bases 11-283) of *S. kambucha wtf4^poison^* coding sequence and this construct was synthesized and cloned into pSZB1460 by IDT to generate pSZB1556.

###### S. kambucha wtf4-ex3Δ

We deleted exon 3 (bases 284-541) of *S. kambucha wtf4^poison^* coding sequence and this construct was synthesized and cloned into pSZB1460 by IDT to generate pSZB1557.

###### S. kambucha wtf4- ex4Δ

We deleted exon 4 (bases 542-733) of *S. kambucha wtf4^poison^* coding sequence and this construct was synthesized and cloned into pSZB1460 by IDT to generate pSZB1558.

###### S. kambucha wtf4-ex5Δ

We deleted exon 5 (bases 734-796) of *S. kambucha wtf4^poison^* coding sequence and this construct was synthesized and cloned into pSZB1460 by IDT to generate pSZB1559.

###### S. kambucha wtf4-ex6Δ

We deleted exon 6 (bases 797-885) of *S. kambucha wtf4^poison^* coding sequence and this construct was synthesized and cloned into pSZB1460 by IDT to generate pSZB1560. To construct the mutant antidote, we deleted exon 6 and this construct was synthesized and cloned into pSZB1537 by IDT to generate pSZB1899.

###### S. kambucha wtf4-consΔ

From previous analysis of *S. pombe wtf* genes, a conserved region within exon 3 was identified [20]. This conserved region within exon 3 was 29 base pairs long (bases 284-312) in *S. kambucha wtf4^poison^* coding sequence. To maintain in-frame codons, we included one base pair upstream the conserved region, and made a 30 base pairs deletion (bases 283-312) in *S. kambucha wtf4^poison^* coding sequence and this construct was synthesized and cloned into pSZB1460 by IDT to generate pSZB1617.

###### S. kambucha wtf4-Δ^10-poison^-mEos

We amplified a 10 amino acid truncated poison from pSZB464 [15] with oligos 3183+3186 and cloned this into V08 [30] via Golden Gate assembly (New England Biolabs) to generate pSZB1402. We digested this with SacI and KpnI and cloned the insert into SacI, KpnI digested pRS316 [70] to generate pSZB1505.

###### S. kambucha wtf4-Δ^10-antidote^-mCherry

We amplified 10 amino acid truncated antidote from pSZB708 [15] with oligos 1402+3138, and mCherry-cyc1T from pSZB708 [15] with oligos 3139+2170. We then stitched these pieces with oligos 1402+2170. We cut this fragment with XhoI, BamHI and cloned this into XhoI, BamHI cut pDK20 [69] to generate pSZB1416. We cut this plasmid with KpnI and XhoI and cloned the insert into KpnI, XhoI cut pRS314 [70] to generate pSZB1550.

###### S. kambucha wtf4-Δ^20-poison^-mEos

We amplified 20 amino acid truncated poison from pSZB464 [15] with oligos 3183+3282 and cloned this into V08 [30] via Golden Gate assembly (New England Biolabs) to generate pSZB1444. We digested this with SacI and KpnI and cloned the insert into SacI, KpnI digested pRS316 [70] to generate pSZB1507.

###### S. kambucha wtf4-Δ^20-antidote^-mCherry

We amplified 20 amino acid truncated antidote from pSZB708 [15] with oligos 1402+3829. We digested this fragment with XhoI and BamHI and cloned the insert into XhoI, BamHI cut pSZB1537 to generate pSZB1567.

###### S. kambucha wtf4^poison^-mCherry

We amplified *S. kambucha wtf4^poison^-mCherry-cyc1T* from pSZB708 [15] with oligos 2625+964. This product was cut with BamHI and XhoI and cloned into BamHI, XhoI cut pDK20 [69] to generate pSZB1374. We cut this plasmid with KpnI and XhoI and cloned the insert into KpnI, XhoI cut pRS314 [70] to generate pSZB1381.

###### S. kambucha wtf4^poison^-mEos

We digested RHX1389 [15] with KpnI, SacI and ligated the insert with KpnI, SacI cut pRS316 [70] to generate pSZB1455. This plasmid had the start site of the poison mutated to TAG, which was then corrected to ATG by GenScript to generate pSZB1476.

###### S. kambucha wtf4^poison^-GBP-mCherry

We added the GFP-binding protein sequence from Addgene plasmid #89068 [74] at the end of *S. kambucha wtf4^poison^* and this construct was synthesized and cloned into pSZB1537 by IDT to generate pSZB1748.

###### S. kambucha NT*-wtf4^poison^-mEos

We added the mutated N-terminal domain (D40K, K65D) from the flagelliform spidroin 1A variant 1 from *Trichonephila clavipes*, NT* [34,35], followed by a TEV cleavage site (ENLYFQS) [75] at the N-terminus of *S. kambucha wtf4^poison^* coding sequence, which was synthesized and cloned into pSZB1460 by IDT to generate pSZB1900.

###### S. kambucha MBP-wtf4^poison^-mEos

We added the *S. cerevisiae* codon-optimized *E. coli* Maltose Binding Protein (MBP) coding sequence [76,77] followed by a 4X(GGGS)-GG linker to the N-terminus of *S. kambucha wtf4^poison^*coding sequence, which was synthesized and cloned into pSZB1460 by IDT to generate pSZB1949.

###### S. kambucha wtf4^antidote^-GFP

We ordered *S. kambucha wtf4^antidote^* which was synthesized and cloned into pSZB1540 by IDT to generate pSZB1874.

###### S. kambucha wtf4^antidote^-mCherry

We amplified *S. kambucha wtf4^antidote^-mCherry-cyc1T* from pSZB1005 [15] with oligos 1402+2170. This product was digested with BamHI and XhoI and cloned into BamHI, XhoI cut pDK20 [69] to generate pSZB1699. This plasmid was digested with KpnI, XhoI and ligated with KpnI, XhoI cut pRS314 [70] to generate pSZB1774.

###### S. kambucha wtf4^antidote^-mEos

We digested pSZB1120 [7] with KpnI, SacI and ligated the insert into KpnI, SacI cut pRS316 [70] to generate pSZB1453. This plasmid had the start site of the antidote mutated to TAG, which was then corrected to ATG by GenScript to generate pSZB1477.

###### S. octosporus wtf25^poison^-mEos

We amplified *S. octosporus wtf25^poison^* from pSZB1353 [19] with oligos 3841+3840 and cloned this into V08 [30] via Golden Gate assembly (New England Biolabs) to generate pSZB1548. We digested this with SacI and KpnI and cloned the insert into SacI, KpnI digested pRS316 [70] to generate pSZB1585.

###### S. octosporus wtf25^poison^-mCherry

We ordered *S. octosporus wtf25^poison^* coding sequence was synthesized and cloned into pSZB1537 by IDT to generate pSZB1807.

###### S. octosporus wtf25^poison^-GBP-mCherry

We added the GFP-binding protein sequence from Addgene plasmid #89068 [74] at the end of *S. octosporus wtf25^poison^* coding sequence and this construct was synthesized and cloned into pSZB1537 by IDT to generate pSZB1868.

###### S. octosporus wtf25^antidote^-mCherry

We ordered *S. octosporus wtf25^antidote^* coding sequence was synthesized and cloned into pSZB1537 by IDT to generate pSZB1746.

###### S. octosporus wtf25^antidote^-mEos

We ordered *S. octosporus wtf25^antidote^* coding sequence was synthesized and cloned into pSZB1460 by IDT to generate pSZB1806.

###### S. octosporus wtf25^antidote^-mEos in pRS314

We amplified *S. octosporus wtf25^antidote^* from pSZB1347 [19] with oligos 3839+3840. This insert was cloned into V08 [30] via Golden Gate cloning (New England Biolabs) to generate pSZB1593. We digested this with KpnI, SacI and cloned the insert into pRS314 [70] to generate pSZB1598.

###### S. octosporus wtf25^antidote^-GFP

We ordered *S. octosporus wtf25^antidote^* coding sequence was synthesized and cloned into pSZB1540 by IDT to generate pSZB1869.

###### S. octosporus wtf25-repΔ

We deleted 30 base pairs (bases 505-534) that make up the exon 4 repeats in *S. octosporus wtf25^poison^* and was synthesized and cloned into pSZB1460 by IDT to generate pSZB1687. To construct the mutant antidote, we deleted bases 640-669 of the antidote coding sequence and this construct was synthesized and cloned it into pSZB1540 by IDT generate pSZB1739.

###### S. octosporus wtf25-rep^Sk^

To swap the coding sequence repeats between *S. kambucha wtf4* exon 6 and *S. octosporus wtf25* exon 4, we replaced the 30 base pairs from *wtf25* (ATAGGAAACGGTGCACGGCATAGGAAAT) with 30 base pairs from *wtf4* (ATAGGGAATATAGGGAGAGCGTTTAGAGGT) in *S. octosporus wtf25*. The mutant poison was synthesized and cloned into pSZB1537 by IDT to generate pSZB1732. The mutant antidote was synthesized and cloned into pSZB1460 by IDT to generate pSZB1731.

###### S. octosporus wtf25Δ^10-poison^-mEos

We amplified a 10 amino acid C-terminal truncated *wtf25^poison^* from pSZB1353 [19] with oligos 4173+4174 and cloned this into V08 [30] via Golden Gate assembly (New England Biolabs) to generate pSZB1675. We then cut out the tagged poison with SacI and KpnI and ligated it with cut pRS316 [70] to generate pSZB1694.

###### S. octosporus NT*-wtf25^poison^

We added the mutated N-terminal domain (D40K, K65D) from the flagelliform spidroin 1A variant 1 from *Trichonephila clavipes*, NT* [33,34], followed by a TEV cleavage site (ENLYFQS) [75] at the N-terminus of *S. octosporus wtf25^poison^* coding sequence, which was synthesized and cloned into pSZB1460 by IDT to generate pSZB1927.

###### S. octosporus MBP-wtf25^poison^-mEos

We added the *S. cerevisiae* codon-optimized *E. coli* Maltose Binding Protein (MBP) coding sequence [76,77] followed by a 4X(GGGS)-GG linker to the N-terminus of *S. octosporus wtf25^poison^* coding sequence, which was synthesized and cloned into pSZB1460 by IDT to generate pSZB1950.

###### S. octosporus wtf61^poison^-mEos

We amplified *S. osmophilus wtf61^poison^* from pSZB1040 [19] with oligos 3837+3838 and cloned this into V08 [30] via Golden Gate assembly (New England Biolabs) to generate pSZB1569. We digested this with SacI and KpnI and cloned the insert into SacI, KpnI digested pRS314 [70] to generate pSZB1583.

###### S. octosporus wtf61^poison^-mCherry

We amplified *S. osmophilus wtf61^poison^* from pSZB1040 [19] with oligos 4193+4194. We digested this product with XhoI, BamHI and cloned this into XhoI, BamHI cut pSZB1537 to generate pSZB1706.

###### S. octosporus wtf61^antidote^-GFP

We amplified *S. osmophilus wtf61^antidote^* from pSZB1095 [19] with oligos 4192+4194. We digested this product with XhoI, BamHI and cloned this into with XhoI, BamHI cut pSZB1540 to generate pSZB1708.

###### S. octosporus wtf61^antidote^-mEos

We amplified *S. osmophilus wtf61^antidote^* from pSZB1095 [19] with oligos 3836+3837 and cloned into V08 [30] via Golden Gate assembly (New England Biolabs) to generate pSZB1645. We then digested pSZB1645 with KpnI and SacI and ligated it with KpnI, BamHI cut pRS314 to generate pSZB1647.

###### S. cryophilus wtf1^poison^-mEos

We amplified *S. cryophilus wtf1^poison^* from pSZB1122 [19] with oligos 3844+3843 and cloned this into V08 [30] via Golden Gate assembly (New England Biolabs) to generate pSZB1544. We digested this with SacI and KpnI and cloned the insert into SacI, KpnI digested pRS316 [70] to generate pSZB1575.

###### S. cryophilus wtf1^poison^-mCherry

We ordered *S. cryophilus wtf1^poison^* which was synthesized and cloned into pSZB1537 by IDT to generate pSZB1870.

###### S. cryophilus wtf1^antidote^-mEos

We amplified *S. cryophilus wtf1^antidote^* from pSZB1192 [19] with oligos 3843+3842 and cloned this into V08 [30] via Golden Gate assembly (New England Biolabs) to generate pSZB1605. We digested this with SacI and KpnI and cloned the insert into SacI, KpnI digested pRS314 [70] to generate pSZB1612.

###### S. cryophilus wtf1^antidote^-GFP

We ordered *S. cryophilus wtf1^antidote^* which was synthesized and cloned into pSZB1540 by IDT to generate pSZB1871.

###### S. cryophilus wtf1Δ^10-poison^

We amplified a 10 amino acid C-terminal truncated *wtf1^poison^*from pSZB1122 [19] and oligos 4177+4178 and cloned this into V08 [30] via Golden Gate assembly (New England Biolabs) to generate pSZB1673. We then cut out the tagged poison with SacI, KpnI and cloned into cut pRS316 [70] to generate pSZB1692.

###### S. osmophilus wtf41^poison^-mEos

We cloned *S. osmophilus wtf41^poison^* from pSZB1327 [19] with oligos 3361+3362 and cloned this into V08 [30] via Golden Gate assembly (New England Biolabs) to generate pSZB1533. We then cut out the tagged poison with SacI, KpnI and cloned into cut pRS316 [70] to generate pSZB1581.

###### S. osmophilus wtf41Δ^10-poison^-mEos

We cloned C-terminus 10 amino acid truncated *S. osmophilus wtf41^poison^* from pSZB1325 [19] with oligos 4175+4176 and cloned this into V08 [30] via Golden Gate assembly (New England Biolabs) to generate pSZB1676. We then cut out the tagged poison with SacI, KpnI and cloned into cut pRS316 [70] to generate pSZB1696.

###### S. osmophilus wtf41^antidote^-mEos

We amplified *S. osmophilus wtf41^antidote^* from pSZB1325 [19] with oligos 3832+3362 and cloned this into V08 [30] via Golden Gate assembly (New England Biolabs) to generate pSZB1599. We then cut out the tagged poison with SacI, KpnI and cloned into cut pRS314 [70] to generate pSZB1607.

###### S. osmophilus wtf19^poison^-mEos

We amplified *S. osmophilus wtf19^poison^* from pSZB1324 [19] with oligos 3835+3834 and cloned this into V08 [30] via Golden Gate assembly (New England Biolabs) to generate pSZB1546. We then cut out the tagged poison with SacI, KpnI and cloned into cut pRS316 [70] to generate pSZB1579.

## Strain construction

### S. cerevisiae strains with Galactose and β-estradiol inducible systems

We used the previously published strain, SZY1637 [15] and an independently constructed isolates of SZY1637, SZY1639 or SZY5807, to construct the strains in this study by transforming in the appropriate vectors. These strains have the lexA-ER-haB42 transcription factor integrated at *HIS3* [78]. In general, we used a protocol, modified from [79], to transform the vectors into these strains. Briefly, we incubated a mixture of 276μL of PLATE solution, 50μL of boiled salmon sperm DNA, 30μL of water, 1-4μL of vector DNA and a match-head sized amount of yeast at 30°C overnight and plated on selective media on the following day to select for transformants. We used Synthetic Complete (SC) media (Yeast Nitrogen Base from VWR #291940 and SC Supplement mix from SunSci) with the appropriate dropouts of the selective components to maintain vectors in the strains as in [15]. In some cases, transformants did not express the construct when induced. To ensure proper testing of genotypes, we screened multiple transformants via cytometry or microscopy to ensure that strains express the fluorescently tagged construct(s).

#### Construction of S. cerevisiae strain with mEosNB-FTH1

We mated RHY3171 [31] to BY4741 [80] to obtain the progeny (SZY6594) that contained the *ho!1::natMX::tTa{Off}tet07^mEosNB-(G4s)3-FTH1* construct, but lacked the *pGal-WHI5::hphMX* construct.

## Spot assays

For spot assays, we first grew the strains for each experiment in the appropriate SC dropout media overnight at 30°C. The next day, we measured the OD600 of each strain and normalized the OD of all the strains to an OD of ∼1. We then serially diluted these normalized cultures 10- fold (up to the dilution of 10^-5^) in water and plated 5μL of each dilution on both SC dropout media and SC Galactose dropout media. We imaged the plates post 72h of growth at 30°C, except in the case of Figs 4B, S6B and S6C, where plates were grown for an additional day at 30°C to clearly visualize differences in poison toxicity.

For the spot assays presented in Fig 3D, we first grew the strains in 5mL SC-URA (uracil) media overnight at 30°C. The next day, we pelleted overnight cultures and resuspended in 1mL of sterile water. We then used 100μL of this as the first in the dilution series. This was diluted 5- fold, up to eleven dilutions in order to observe the toxicity/suppression clearly. We plated 5μL of each dilution on SC-URA media, SC Galactose-URA media and SC Galactose-URA media with 40mg/L Doxycycline (Calbiochem #324385). We imaged the plates post 72h of growth at 30°C.

For the spot assays presented in S7B Fig, we first grew the strains in 5mL SC-TRP-URA media overnight at 30°C. The next day, we pelleted overnight cultures and resuspended in 1mL of sterile water. We then used 100μL of this as the first in the dilution series. This was diluted 5- fold, up to 11 dilutions in order to observe the toxicity clearly. We plated 5μL of each dilution on SC-TRP-URA media and SC Galactose-TRP-URA media. We imaged the plates post 72h of growth at 30°C.

## Microscopy

The appropriate strains for each experiment were first grown in 5mL of the appropriate SC dropout media overnight at 30°C. The next day, we inoculated 3mL of SC Raffinose dropout media with 1mL of the saturated overnight cultures and let them grow overnight at 30°C. On the following day, we pelleted the cultures and resuspended the cells in 3mL SC Galactose dropout media and incubated at 30°C. Cells were induced for 4-6 hours on SC Galactose dropout media and then imaged on a Zeiss LSM 980 confocal microscope which consisted of an Axio Observer Z1 base, through a 40x C-Apochromat (NA = 1.2) water immersion objective. The GFP and mCherry tagged proteins were excited at 488 and 561 nm, respectively. Emission was collected in channel mode onto a GaAsP detector in photon counting mode through a 491-544 nm bandpass for GFP and a 570-632 nm bandpass for mCherry. Transmitted light was also collected. Image fields of view were zoomed optically to a ∼26 µm square with 512 pixels in each dimension.

For imaging cells expressing *S. octosporus* Wtf61^antidote^-GFP, *S. kambucha* Wtf4^poison^-GBP-mCh and Rnq1-mCardinal, we first grew cells in SC-Trp-Ura-Leu media. The next day, we inoculated 3mL of SC Raffinose-Trp-Ura-Leu media with 1mL of the saturated overnight cultures and let them grow overnight at 30°C. The following day, we pelleted the cultures and resuspended the cells in 3mL SC Galactose-Trp-Ura-Leu + 500nM β-estradiol (VWR, # #AAAL03801-03) media and incubated at 30°C. Cells were induced for 4 hours and then imaged as described above, except that we also excited mCardinal tagged proteins at 640nm and collected the emission on channel mode onto a 650-700 nm bandpass detector of the Zeiss LSM 980 confocal microscope.

## Spectral unmixing for mEosNB-FTH1 strains

For imaging the mEosNB-FTH1 strains, we first grew the strains in 5mL SC-URA+40mg/L Doxycycline. The next day, we inoculated 3mL of both SC Raffinose-URA and SC Raffinose- URA+40mg/L Doxycycline media with 1mL of the saturated overnight cultures and let them grow overnight at 30°C. The following day, we pelleted the cultures and resuspended the cells in 3mL of both SC Galactose-URA and SC Galactose-URA+40mg/L Doxycycline and incubated at 30°C. The rest of the methodology was as described in the section above.

The background fluorescence for strains constructed with SZY6594 had high autofluorescence in the GFP channel. To distinguish autofluorescence from true mEos signal, we spectrally unmixed the images. First, we captured spectral images with the lambda mode on the detector of the Zeiss LSM 980 confocal microscope (same settings as above), exciting the cells at 488 nm and collecting emission over the entire visual spectrum. We then made reference spectra for mEos and autofluorescence from control cells, and then spectrally unmixed images using an in-house written plugin on ImageJ (https://research.stowers.org/imagejplugins/). The data presented in Fig 3C has transmitted light images collected on the channel mode with GFP excitation, and the spectrally unmixed images collected on lambda mode.

## Image analysis to quantify colocalization of mEos and mCherry signals in tagged constructs

Cell regions of interest (ROIs) were found based on the transmitted light only using Cellpose (https://www.cellpose.org/) in Python using the “cyto” model and a diameter of 200. Pixels from these ROIs were then background subtracted using an average background signal from an ROI placed away from any cell. The Scipy Pearson Correlation for each cell ROI was then calculated for all pixels in each cell. Dead cells and aberrant ROIs were hand filtered out using Fiji (https://fiji.sc/). Find the raw data for these analyses in S1 Data.

## DAmFRET

We performed the analysis similar to the methods in [15], with a few exceptions. We grew the strains in the same method described for microscopy, however, we induced the cells for 5-6 hours. We then photoconverted the cells post induction for 5 minutes, using the OmniCure S2000 Elite UV lamp for photoconversion. The total power over 5 minutes of exposure amounts to 12.048 J/cm^2^. The sample collection and data analysis were similar to methods described in [15] with a few exceptions. We included an empty vector strain in each experiment to effectively exclude auto fluorescent cell populations. All the cells for each experiment were induced at the same time, for the same amount of time, and were grown in the same 96-well plate. Three technical replicates were included for each sample.

There was only one fluorescent population that was independent of expression level for each construct analyzed. In order to clearly visualize AmFRET values across the cells expressing different constructs, we analyzed cells above a certain Acceptor fluorescence intensity threshold set for each experiment. For each set of experimental data, the median of the Wtf4^poison^-mEos AmFRET values was calculated. The difference of this median value from 1 was then added to all the datasets within an experiment. For visualizing the results of the DAmFRET experiments, plots of AmFRET values were generated using GraphPad Prism 10 (Version 10.2.3 (347)).

Statistical analysis for the DAmFRET experiments involved pooling the technical replicates and performing pairwise t-tests comparing these values to either the wild-type Wtf4^poison^-mEos dataset, or the monomer dataset. For each DAmFRET experiment, we then did a multiple sampling correction (Bonferroni correction) by dividing the p value cutoff (0.05) by the number of tests performed. Find the raw data and statistical analysis for each DAmFRET experiment in S4 Data.

We used GraphPad Prism 10 (Version 10.2.3 (347)) to visualize acceptor fluorescence intensity values of live cells expressing the relevant Wtf^poison^ constructs presented in S6 Fig. Three technical replicates of each isolate were plotted, with the median included. Find the raw data for this analysis in S2 Data.

## Acknowledgements

We thank the members of the Zanders lab, the Halfmann lab and the Core facilities at Stowers Institute for their help and guidance throughout the stages of development of this paper.

---

**S1 Figure.**
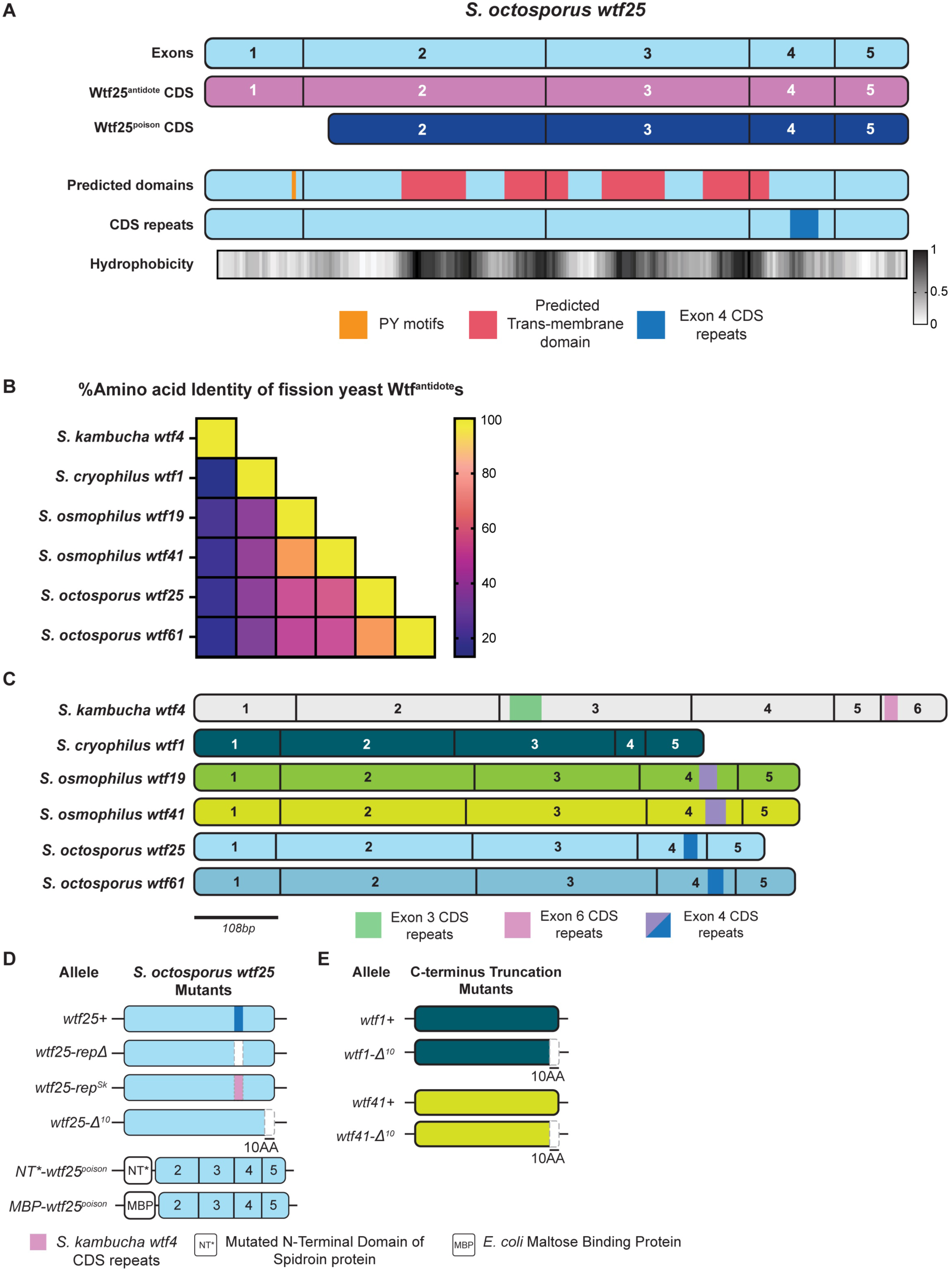
Wtf proteins share limited amino acid identity but have common features **A.** A cartoon of *S. octosporus wtf25* coding sequence (CDS). Wtf25^antidote^ coding sequence is shown in purple, which includes exons 1-5. The Wtf4^poison^ coding sequence is shown in navy, which begins at the 27th base pair of exon 2 and extends through exon 5. Row 4 depicts the predicted transmembrane domains (in red) and PY motif (in mustard). Row 5 depicts the CDS repeats found in exon 4 (in cobalt). Row 6 depicts the normalized hydrophobicity of Wtf25 proteins from ProtScale, with the Kyle and Doolittle Hydropathy scale [81]. The higher the number on the scale, the higher the hydrophobicity of the amino acid. See S2 and S3 Tables for more detailed descriptions. **B.** Pairwise amino acid identity of the 6 Wtf^antidote^ proteins shown. The amino acid sequences were aligned using Geneious Prime (2023.0.4), and the percentage amino acid identity is depicted as a heatmap, with yellow being 100% identity. **C.** Depiction of CDS repeats and lengths of 6 *wtf^antidote^* CDSs. The CDS repeats found in exon 6 of *S. kambucha wtf4* are homologous to those found in exon 4 of the other *wtf* genes [19]. The scale bar represents 108 base pairs (bp). **D-E.** *S. octosporus wtf25* (**D**), *S. cryophilus wtf1*, and *S. osmophilus wtf41* (**E**) mutants constructed in this study. The categories have their respective wild type allele shown on top. See S1 Table for a comprehensive overview of the alleles and their phenotypes.

**S2 Figure.**
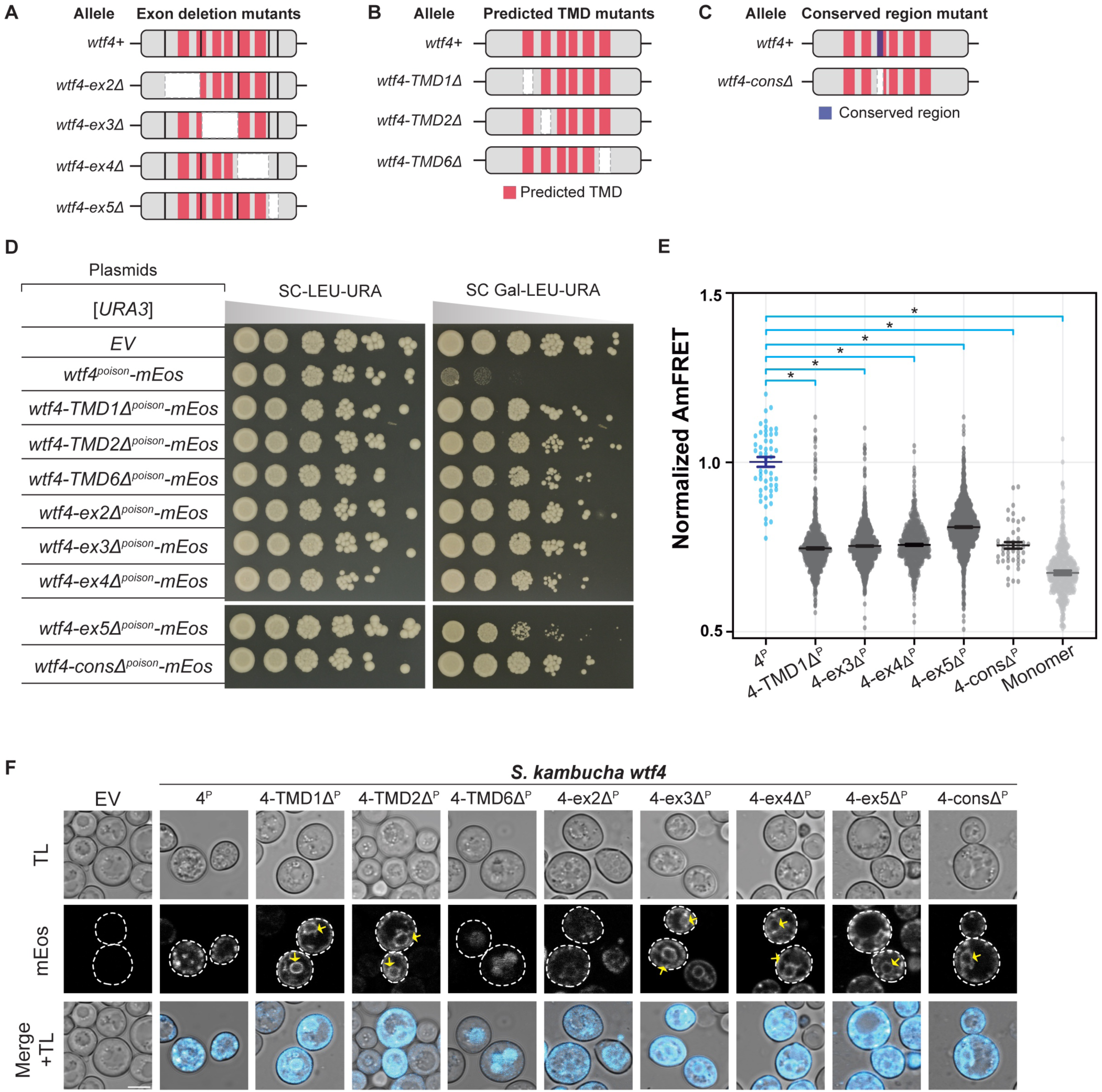
Deletion mutants affect Wtf4^poison^ toxicity, self-assembly and localization. Cartoon of *S. kambucha wtf4* exon deletion mutants (**A**), predicted transmembrane domain (TMD) deletion mutants (**B**) and a mutant that deletes a 9 amino acid conserved region encoded in exon 3 (**C**). **D.** A spot assay of cells serially diluted and plated on SC-LEU-URA and SC Gal- LEU-URA plates and grown at 30^·^C for 3 days. Each strain carries an empty [*LEU2*] plasmid, and either an empty [*URA3*] plasmid (EV) or the indicated *wtf4^poison^-mEos* alleles under the control of a galactose-inducible promoter. The horizontal break in the image for each plate is due to rearrangement of the image to facilitate easy comparison. All strains were grown on the same plates (i.e., one SC-LEU-URA or SC Gal-LEU-URA plate). **E.** AmFRET values for three technical replicates of the specified Wtf4^poison^-mEos proteins and monomer-mEos (negative control). The median is indicated with a solid line and the bars represent the interquartile range. For easier comparison, the values were normalized so that Wtf4^poison^ had a median of 1 in each experiment. The data shown here do not include outliers. See S2 Data for the complete dataset and p-values. Statistical significance: *p<0.008, **p<0.0008, t-tests between the means of each replicate, with Bonferroni correction. **F.** Representative images of the same strains depicted in D were induced in galactose media for 4 hours at 30^·^C to express the indicated mEos-tagged proteins. The images are not at the same brightness and contrast settings to clearly show localization of tagged proteins. Yellow arrows indicate endoplasmic reticulum-like localization. 4^P^ indicates Wtf4^poison^, TL is transmitted light, and the scale bar is 4 µm.

**S3 Figure.**
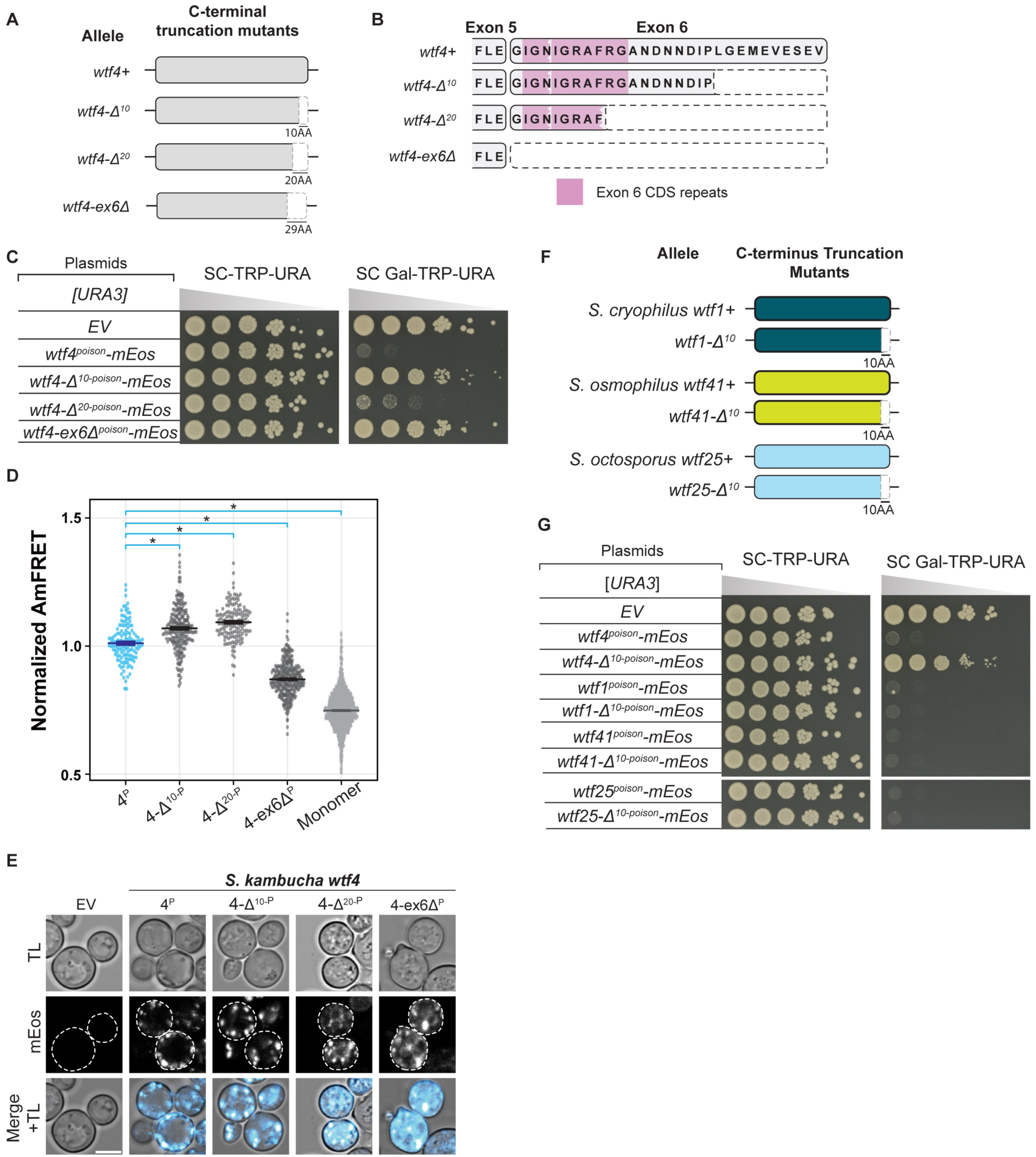
C-terminal truncations have inconsistent effects on Wtf^poison^ protein toxicity. **A.** Cartoon of C-terminal truncation mutants of *S. kambucha wtf4*. **B.** Cartoon illustrating the amino acids lost with the C-terminal truncation alleles of *wtf4*. The amino acids comprising the exon 6 coding sequence repeats are highlighted in pink. **C.** A spot assay of cells serially diluted and plated on SC-TRP-URA and SC Gal-TRP-URA plates and grown at 30^·^C for 3 days. Each strain carries both an empty [*TRP1*] plasmid (EV) and a [*URA3*] plasmid that is either empty or carries the indicated *wtf4^poison^-mEos* allele under the control of a galactose inducible promoter. **D.** AmFRET values for three technical replicates of the specified Wtf4^poison^-mEos alleles and monomer-mEos (negative control). The median is indicated with a solid line and the bars represent the interquartile range. For easier comparison, the values were normalized so that Wtf4^poison^ had a median of 1 in each experiment. The data shown here do not include outliers. See S2 Data for the complete dataset and p-values. Statistical significance: *p<0.0125, t-test with Bonferroni correction. **E.** Representative images of the same strains depicted in C were induced with galactose media for 4 hours at 30^·^C to express the indicated mEos-tagged proteins. The images are not at the same brightness and contrast settings to clearly show localization of tagged proteins. 4^P^ indicates Wtf4^poison^, TL is transmitted light, and the scale bar is 4 µm. **F.** Cartoons of C-terminal truncation mutants of *S. cryophilus wtf1* (*wtf1-!1^10^*)*, S. osmophilus wtf41* (*wtf41-Δ^10^*), and *S. octosporus wtf25* (*wtf25-!1^10^*). **G.** Spot assays of cells serially diluted and plated on SC-TRP-URA and SC Gal-TRP-URA plates and grown at 30^·^C for 3 days. Each strain carries both an empty [*TRP1*] plasmid (EV) and an empty [*URA3*] plasmid (EV), or the indicated *wtf^poison^-mEos* allele under the control of a galactose-inducible promoter. The horizontal breaks in the images for each plate are due to rearrangements of the image to facilitate easy comparison. All strains were grown on the same plates (i.e., one SC-TRP-URA or SC Gal-TRP-URA plate).

**S4 Figure.**
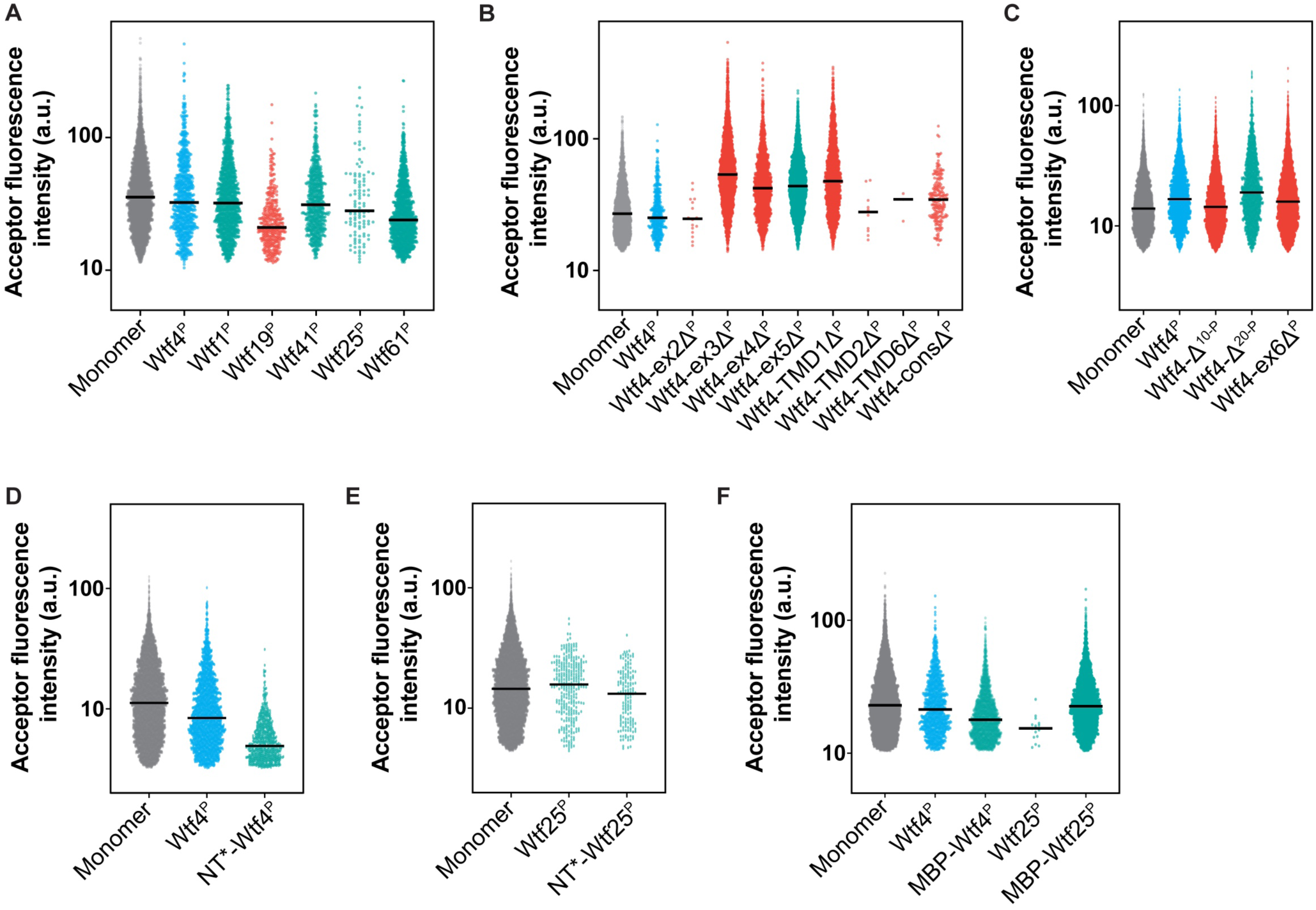
The fluorescence signal intensity of expressed Wtf^poison^ proteins does not correlate with toxicity. **A-F** Acceptor fluorescence intensity (in arbitrary units, a.u.) of similarly sized live cells across all Wtf DAmFRET experiments in this study, with the line indicating the median of the population. Data represented here is from the following experiments: A corresponds to Fig 2B, B corresponds to S2E Fig, C corresponds to S3D Fig, D corresponds to Fig 4C, E corresponds alleles in S6 Fig, and F corresponds to S7D Fig. The Y-axis is scaled to a log10 scale. The data are color coded, with toxic poisons in teal, and non-toxic alleles in orange.

**S5 Figure.**
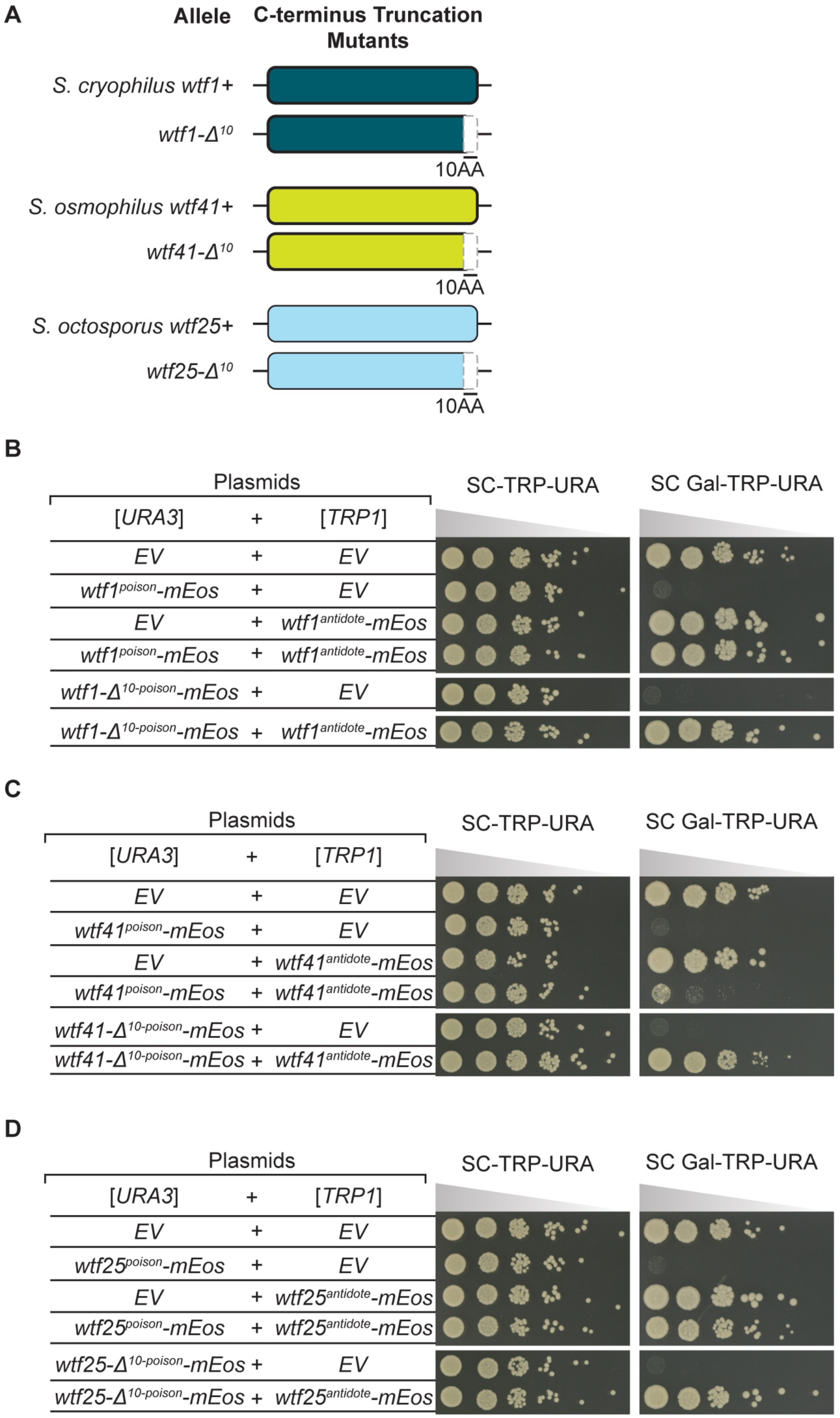
C-terminus truncation alleles of diverged Wtf^poison^s are rescued by their corresponding wild type Wtf^antidote^s. **A.** Cartoon of *S. cryophilus wtf1*, *S. osmophilus wtf41*, and *S. octosporus wtf25* C-terminal truncation mutants. **B-D.** Spot assays of cells serially diluted on SC-TRP-URA and SC Gal-TRP-URA plates and grown at 30^·^C for 3 days. Each strain carries both a [*URA3*] and a [*TRP1*] plasmid. The plasmids are either empty (EV) or carry the indicated *wtf* alleles under the control of galactose-inducible promoters. The horizontal breaks in the images within a panel are due to rearrangements of the images to facilitate easy comparison. All strains within a panel were grown on the same plates (i.e., one SC-TRP-URA or SC Gal-TRP-URA plate).

**S6 Figure.**
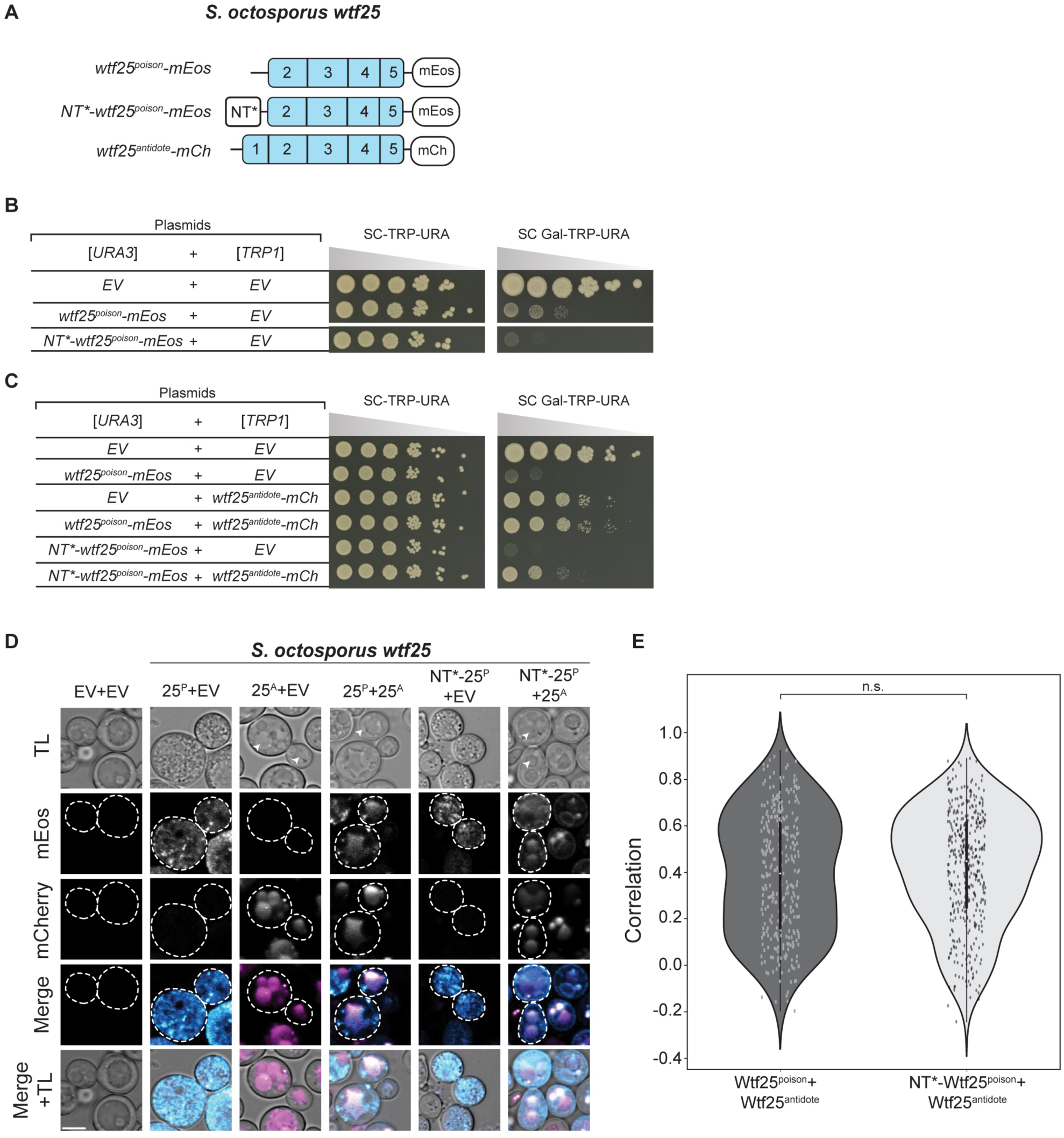
Reducing assembly with NT* tag increases Wtf25^poison^ toxicity. **A.** Cartoon of alleles used in this experiment (B-E). The NT* tag has a general anti-aggregation property [34]. **B-C.** Spot assays of cells serially diluted on SC-TRP-URA and SC Gal-TRP-URA plates and grown at 30^·^C for 4 days. Each strain carries both a [*URA3*] and a [*TRP1*] plasmid. The plasmids are either empty (EV) or carry the indicated *wtf25* alleles under the control of galactose-inducible promoters. In B, the horizontal break in the image of each plate is due to a rearrangement of the image to facilitate easy comparison. All strains within a panel were grown on the same plates (i.e., one SC-TRP-URA or SC Gal-TRP-URA plate for panel B). **D.** Representative images of the same strains shown in C were induced with galactose media for 4 hours at 30^·^C to express the indicated Wtf25^poison^-mEos proteins and/or Wtf25^antidote^-mCherry. The images are not at the same brightness and contrast settings to clearly show localization of tagged proteins. 25^P^ indicates Wtf25^poison^, 25^A^ indicates Wtf25^antidote^, TL is transmitted light, and the scale bar is 4 µm. **E**. Pearson’s Correlation between mCherry and mEos signal in cells from D expressing the specified proteins. N>100, p>0.05, t-test.

**S7 Figure.**
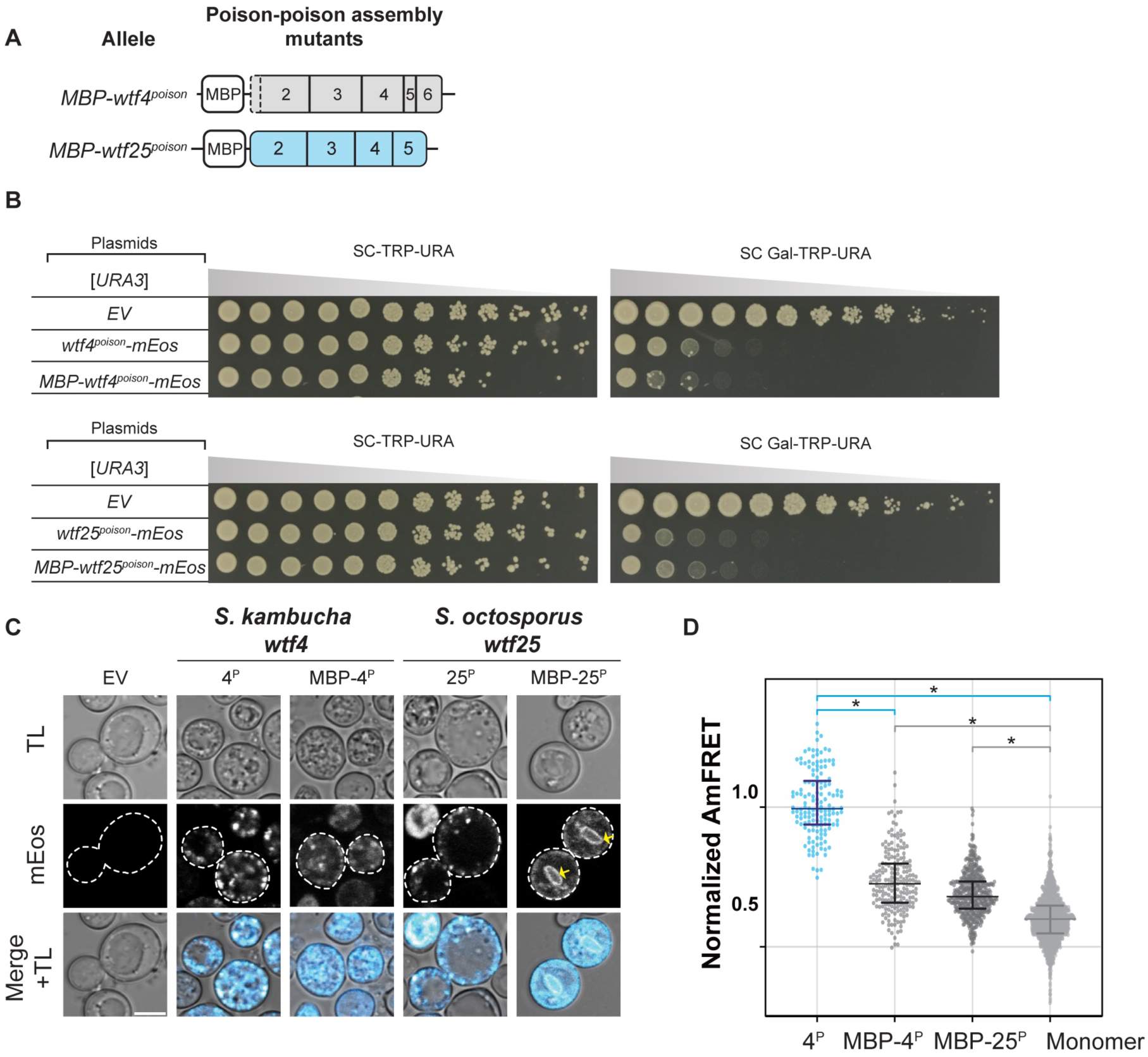
Reducing Wtf^poison^ assembly with MBP tag affects toxicity. **A.** Cartoon of alleles used in this experiment (B-C). MBP is the *E. coli* <u>M</u>altose <u>B</u>inding <u>P</u>rotein [76]. **B.** Spot assay of cells serially diluted and plated on SC-TRP-URA and SC Gal-TRP-URA plates and grown at 30^·^C for 4 days. Each strain carries both an empty [*TRP1*] plasmid and a [*URA3*] plasmid that is either empty (EV) or carries the indicated *wtf^poison^*-*mEos* allele under the control of a galactose-inducible promoter. The horizontal break in the image of each plate is due to rearrangement of the image to facilitate easy comparison. All strains were grown on the same plates (i.e., one SC Gal-TRP-URA plate or SC-Trp-Ura). **C.** Representative images of the same strains depicted in B induced with galactose media for 4 hours at 30^·^C to express the indicated mEos-tagged proteins. The images are not at the same brightness and contrast settings to clearly show localization of tagged proteins. Yellow arrows indicate endoplasmic reticulum-like localization. 4^P^ indicates Wtf4^poison^, 25^P^ indicates Wtf25^poison^, TL is transmitted light, and the scale bar is 4 µm. **D.** AmFRET values for three technical replicates of the specified Wtf^poison^- mEos alleles and monomer-mEos (negative control). The median is indicated with a solid line and the bars represent the interquartile range. For easier comparison, the values were normalized so that Wtf4^poison^ had a median of 1 in each experiment. The data shown do not include outliers. See S2 Data for the complete dataset and p-values. Statistical significance: *p<0.016, t-test with Bonferroni correction.

**S8 Figure.**
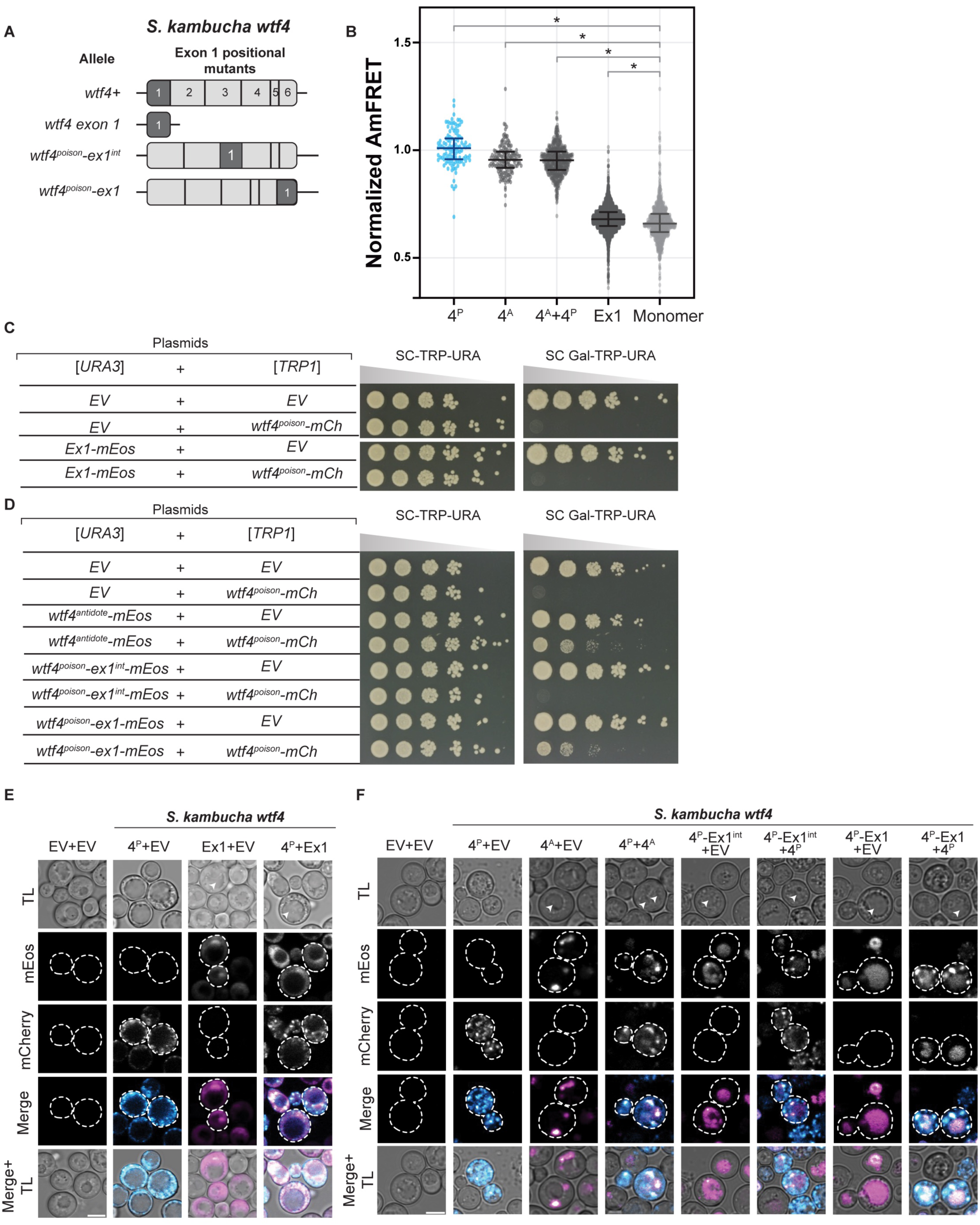
Limited modularity of the Wtf4^antidote^-specific domain. **A.** Cartoon of *wtf4^antidote^* exon 1 and mutants that relocate, or both relocate and mutate the exon. **B.** AmFRET values for three technical replicates of the specified mEos-tagged proteins and monomer-mEos (negative control). The median is indicated with a solid line and the bars represent 1.5 times the interquartile range. For easier comparison, the values were normalized so that Wtf4^poison^ had a median of 1 in each experiment. The data shown do not include outliers. See S2 Data for the complete dataset and p-values. Statistical significance: *p<0.0125, t-tests with Bonferroni correction. **C-D.** Spot assays of cells serially diluted on SC-TRP-URA and SC Gal-TRP-URA plates and grown at 30^·^C for 3 days. Each strain carries both a [*URA3*] and a [*TRP1*] plasmid. The plasmids are either empty (EV) or carry the indicated *wtf4* alleles under the control of galactose-inducible promoters. The horizontal breaks in the images of each plate in panels C and D are due to rearrangements of the images to facilitate easy comparison. All strains within a panel were grown on the same plates (i.e., one SC-TRP-URA or SC Gal-TRP-URA plate for panel C). **E.** Representative images of the same strains shown in C induced with galactose media for 4 hours at 30^·^C to express the Wtf4^poison^-mEos, Wtf4 exon1-mCherry, or both proteins. **F.** Representative images of the same strains shown in D were induced with galactose media for 4 hours at 30^·^C to the indicated *wtf4* alleles. In E-F, arrows in the transmitted light images indicate vacuoles. The images are not at the same brightness and contrast settings to clearly show localization of the tagged proteins. 4^P^ indicates Wtf4^poison^, 4^A^ indicates Wtf4^antidote^, Ex1 is Wtf4 exon1, TL is transmitted light, and the scale bar is 4 µm.

**S9 Figure.**
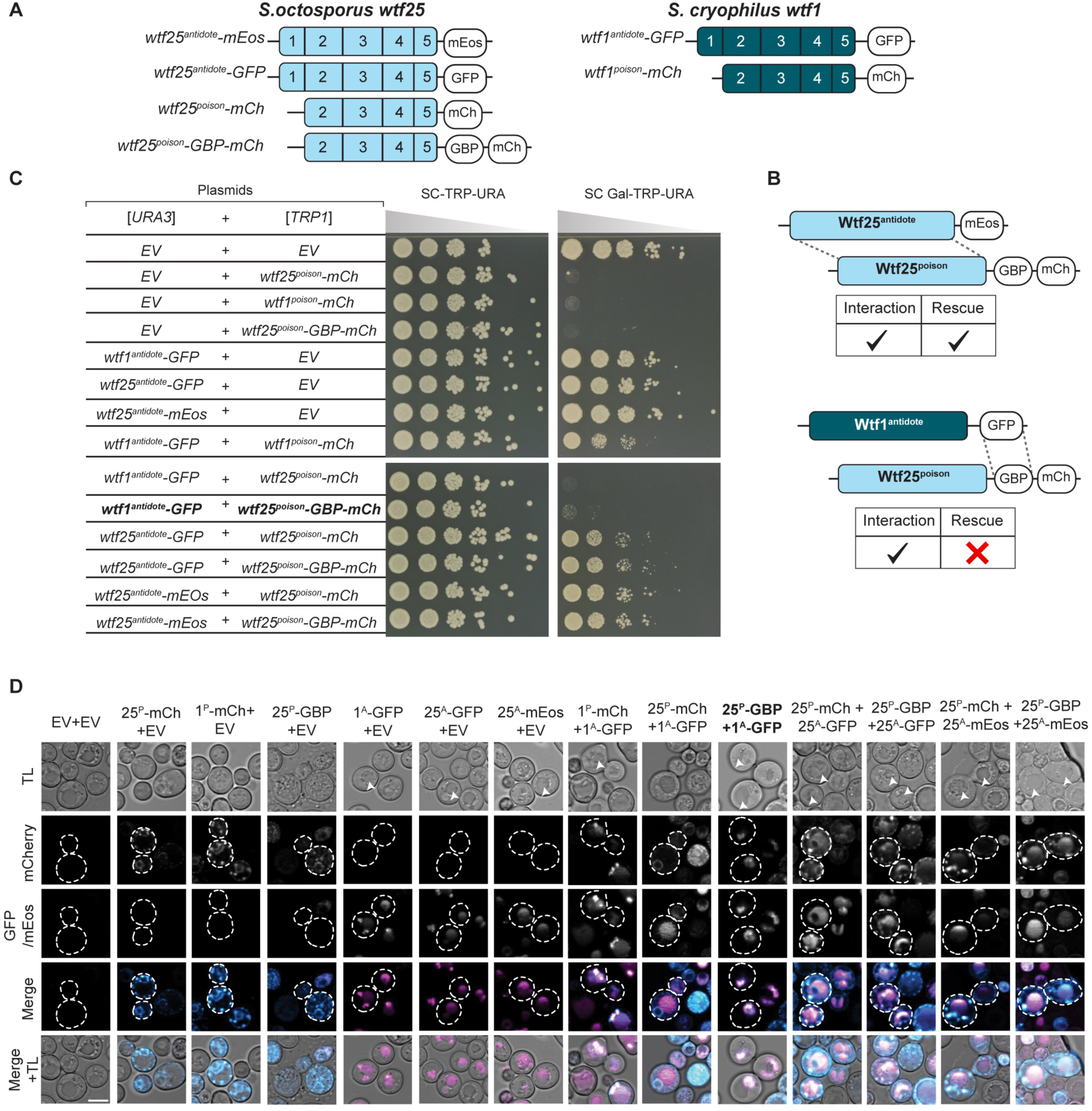
Effective neutralization of Wtf25^poison^ requires more than a physical connection to a Wtf^antidote^_._ **A.** Cartoon of constructs used in this experiment. *S. octosporus* Wtf25^poison^ was tagged at the C-terminus with either mCherry or GBP-mCherry (GBP: GFP-binding protein). *S. octosporus* Wtf25^antidote^ was tagged with mEos or GFP. *S. cryophilus* Wtf1^antidote^ was tagged with GFP. *S. cryophilus* Wtf1^poison^ was tagged with mCherry. **B.** Experimental set up and summary of the results shown in C and D. In a matching Wtf protein pair (top), poison-antidote interaction and rescue of poison toxicity is observed. In the mismatched pair (bottom), interaction between GFP and GBP results in a forced interaction between the poison and the antidote (shown in D). This interaction is insufficient to rescue the mismatched poison (shown in C). **C.** Spot assay of cells serially diluted and plated on SC-TRP-URA and SC Gal-TRP-URA plates and grown at 30^·^C for 3 days. Each strain carries both a [*URA3*] and a [*TRP1*] plasmid. The plasmids are either empty (EV) or carry the indicated *wtf* alleles under the control of galactose-inducible promoters. The horizontal breaks in the images of each plate are due to rearrangements of the image to facilitate easy comparison. All strains within a panel were grown on the same plates (i.e., one SC-TRP-URA or SC Gal-TRP-URA plate). **D.** Representative images the strains depicted in C were induced in galactose for 4 hours at 30^·^C to express the indicated Wtf proteins. The images are not at the same brightness and contrast settings to clearly show localization of tagged proteins. The arrows in the transmitted light images indicate vacuoles. 25^P^ indicates Wtf25^poison^, 25^A^- indicates Wtf25^antidote^, 1^P^ indicates Wtf1^poison^, 1^A^ indicates Wtf1^antidote^, TL indicates transmitted light, and the scale bar is 4 µm.

**S10 Figure.**
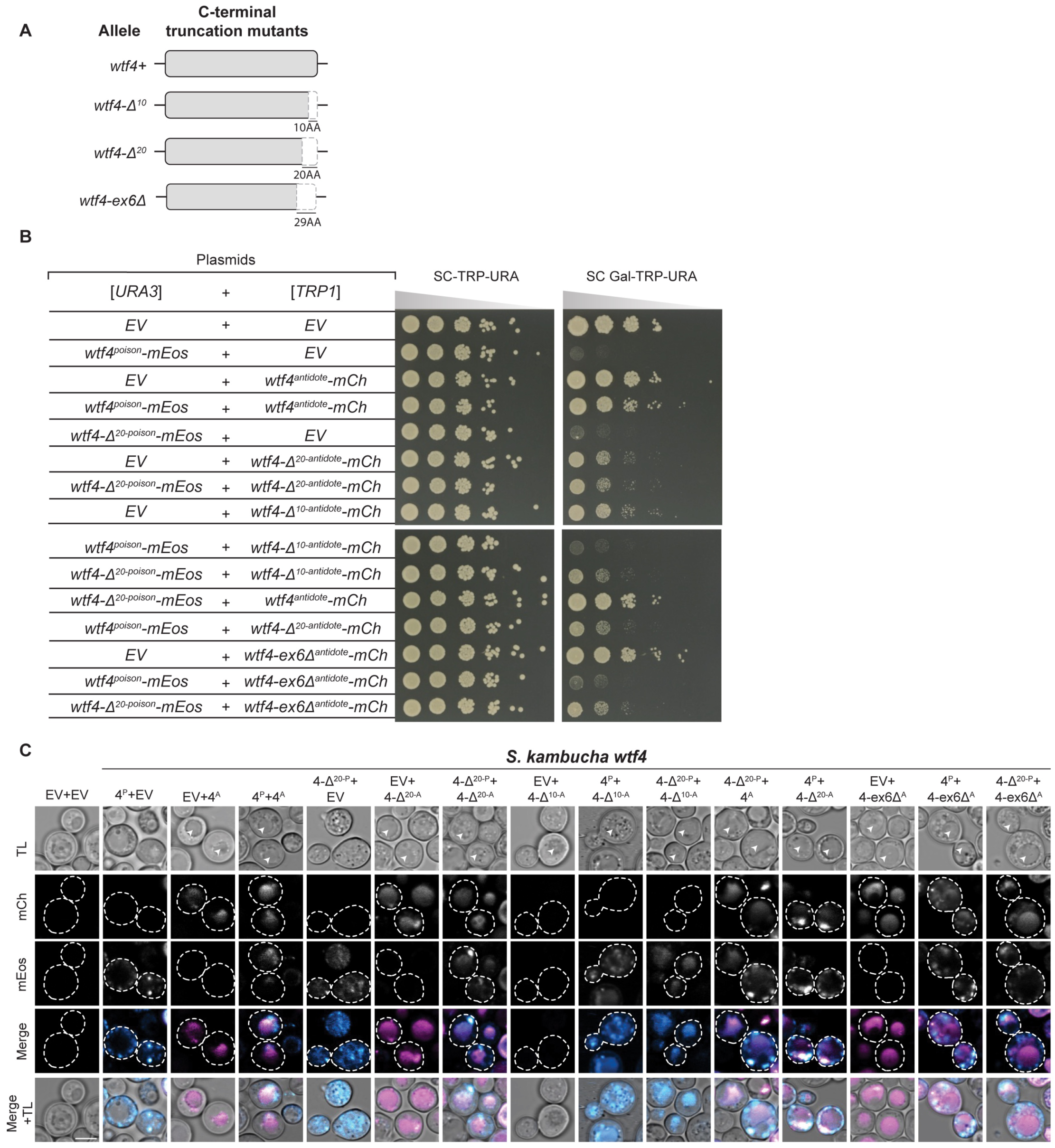
Idiosyncratic poison and antidote compatibility between wild-type and truncation alleles of *wtf4*. **A.** Cartoon of *S. kambucha wtf4* C-terminal truncation mutants. **B.** Spot assay of cells serially diluted and plated on SC-TRP-URA and SC Gal-TRP-URA plates and grown at 30^·^C for 3 days. Each strain carries both a [*URA3*] and a [*TRP1*] plasmid. The plasmids are either empty (EV) or carry the indicated *wtf4* alleles under the control of galactose-inducible promoters. The horizontal breaks in the images are due to rearrangements of the images to facilitate easy comparison. All strains within a panel were grown on the same plates (i.e., one SC-TRP-URA or SC Gal-TRP-URA plate). **C.** Representative images the strains depicted in B were induced in galactose for 4 hours at 30^·^C to express the indicated Wtf4 proteins. The images are not at the same brightness and contrast settings to clearly show localization of tagged proteins. The arrows in the TL panels highlight vacuoles. 4^P^ indicates Wtf4^poison^, 4^A^ indicates Wtf4^antidote^, TL indicates transmitted light, and the scale bar is 4 µm.

**S11 Figure.**
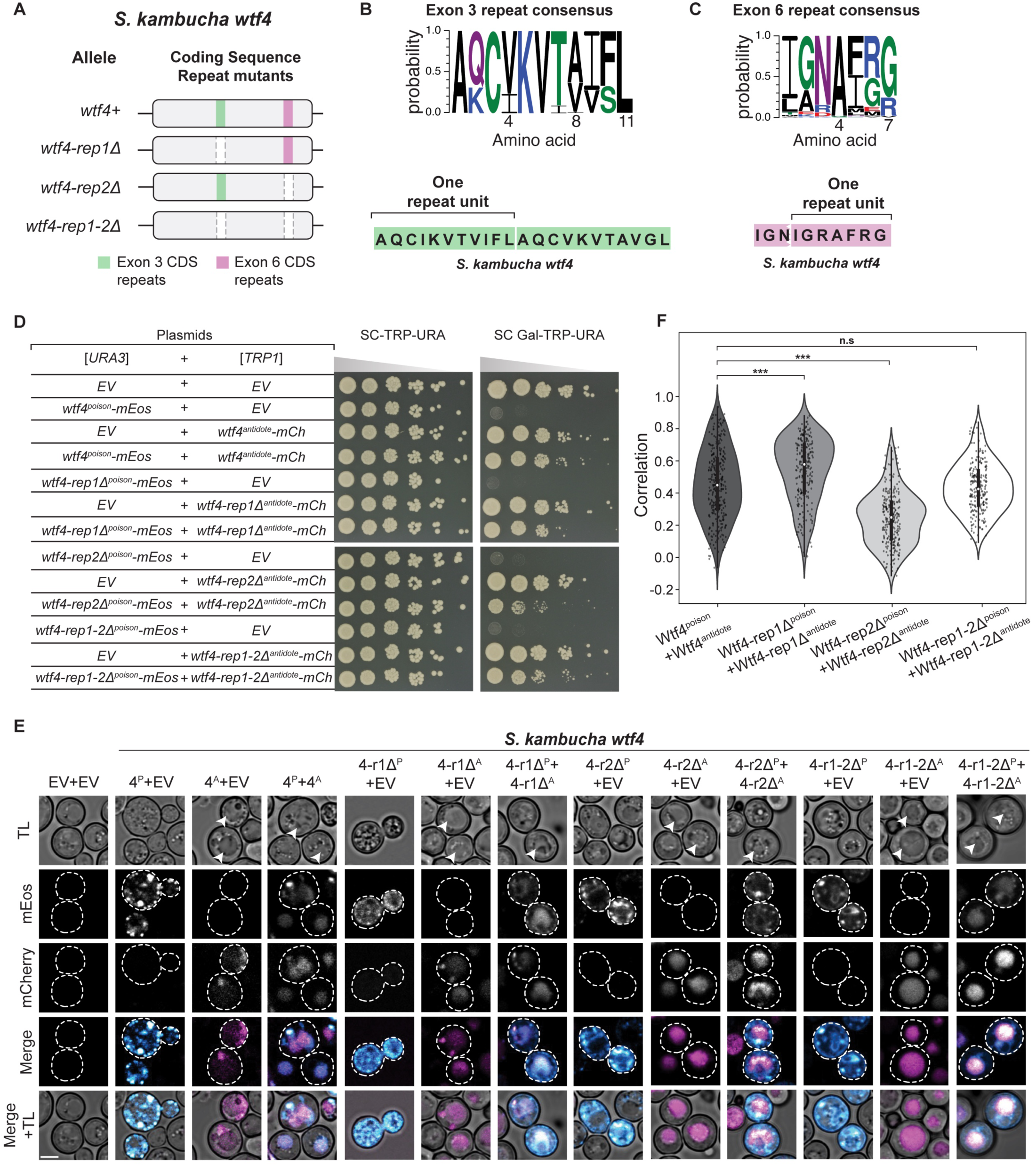
*wtf4* coding sequence repeats are dispensable, but can affect antidote rescue. **A.** Cartoon of exon 3 and exon 6 coding sequence repeat deletion mutants of *S. kambucha wtf4*. **B-C.** Logo for amino acids encoded by the repeats found in exon 3 (B) and exon 6 (C) of *S. pombe wtf* genes from [20]. The sequence of the amino acids encoded in each repeat region in *S. kambucha wtf4* is shown below each logo. **D.** Spot assay of cells serially diluted and plated on SC-TRP-URA and SC Gal-TRP-URA plates and grown at 30^·^C for 3 days. Each strain carries both a [*URA3*] and a [*TRP1*] plasmid. The plasmids are either empty (EV) or carry the indicated *wtf4* alleles under the control of galactose-inducible promoters. The horizontal break in the image of each plate is due to rearrangements of the images to facilitate easy comparison. All strains within a panel were grown on the same plates (i.e., one SC-TRP-URA or SC Gal-TRP-URA plate). **E.** Representative images of the same strains as depicted in D were induced in galactose for 4 hours at 30^·^C to express the indicated Wtf4 proteins. The images are not at the same brightness and contrast settings to clearly show localization of tagged proteins. The arrows in the TL panels highlight vacuoles. 4^P^ indicates Wtf4^poison^, 4^A^ indicates Wtf4^antidote^, TL indicates transmitted light, and the scale bar is 4 µm. **F**. Pearson’s Correlation between mEos and mCherry signal in cells expressing the specified constructs from E. N>100, ***p<0.001, t-test.

**S12 Figure.**
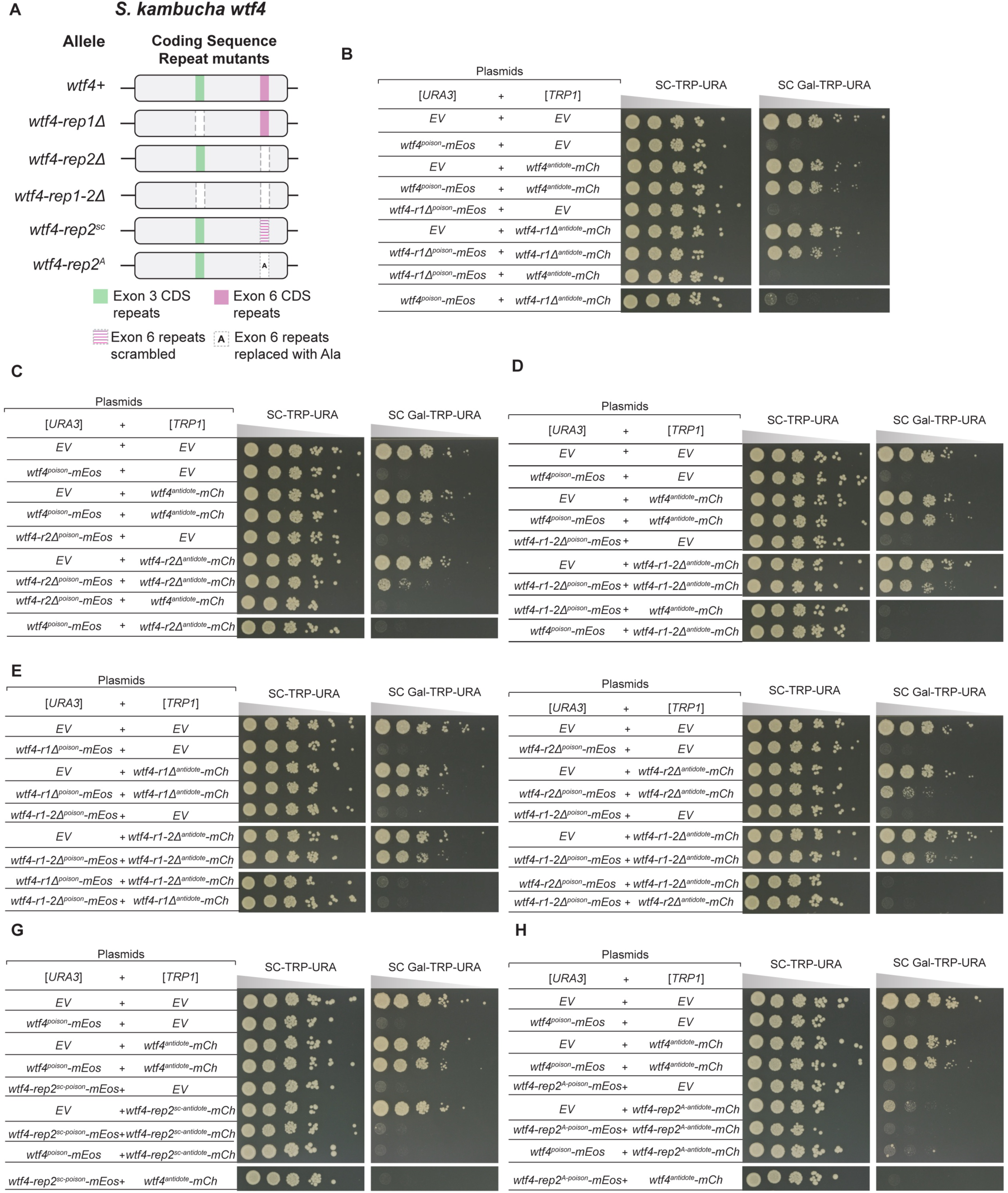
Coding sequence repeat mutant proteins do not interact with wildtype Wtf4 proteins functionally. **A.** Cartoon of coding sequence repeat mutants of *S. kambucha wtf4.* **B-H.** Spot assays of cells serially diluted on SC-TRP-URA and SC Gal-TRP-URA plates and grown at 30^·^C for 3 days. Each strain carries both a [*URA3*] and a [*TRP1*] plasmid. The plasmids are either empty (EV) or carry the indicated *wtf4* alleles under the control of galactose-inducible promoters. The horizontal breaks in the images in within a panel are due to rearrangements of the images to facilitate easy comparison. All strains within a panel were grown on the same plates (i.e., one SC-TRP-URA or SC Gal-TRP-URA plate in panel B).

**S13 Figure.**
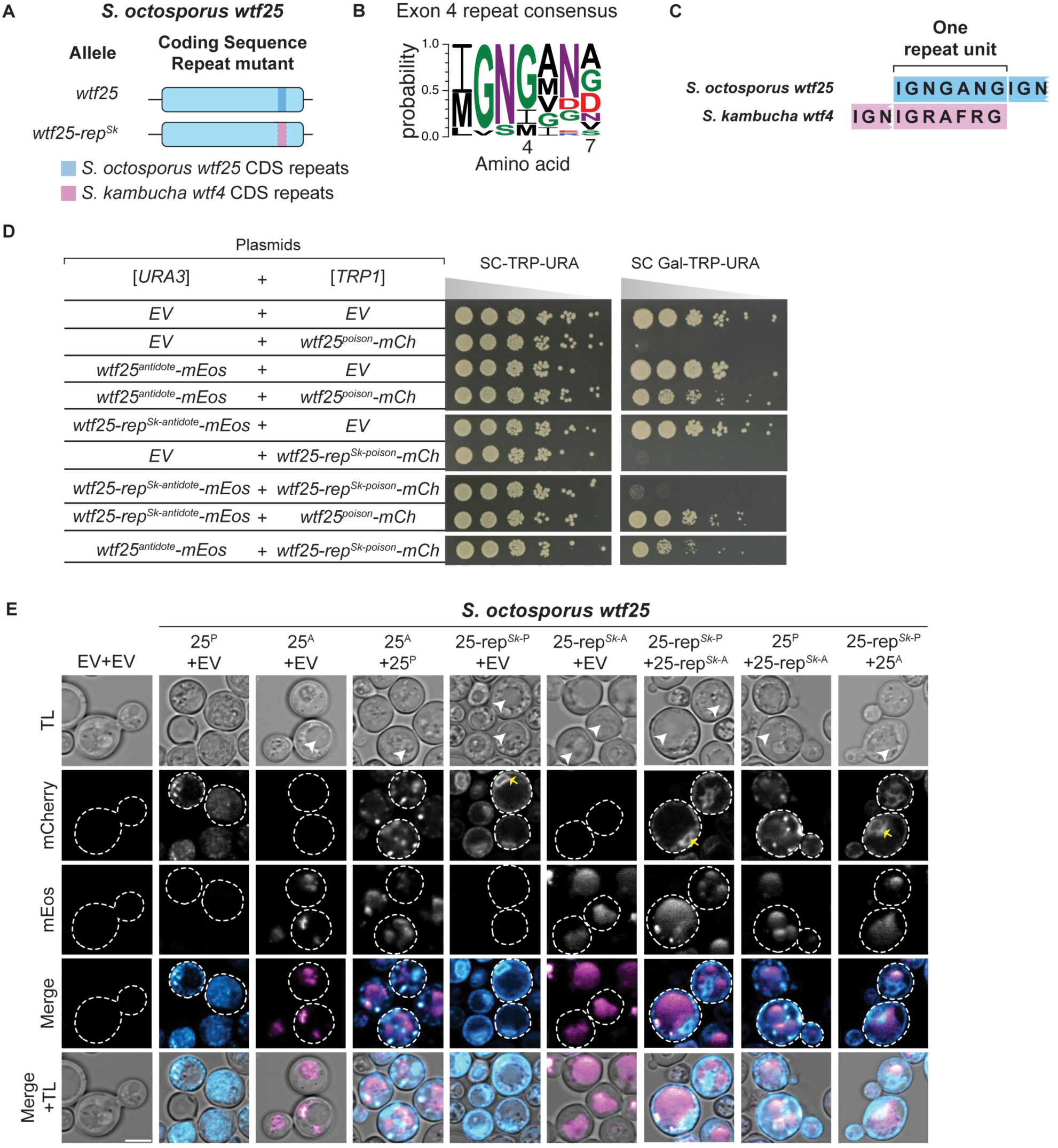
Swapping *wtf* CDS repeats across species affects Wtf25 antidote rescue. **A.** Cartoon of a coding sequence repeat mutant of *S. octosporus wtf25*. **B**. Logo representing the amino acids encoded by the repeats found in exon 4 of *S. octosporus wtf* genes from [19]. **C.** The amino acids encoded by the exon 4 repeats of *S. octosporus wtf25* and by the exon 6 repeats of *S. kambucha wtf4*. **D.** Spot assay of cells serially diluted and plated on SC-TRP-URA and SC Gal-TRP-URA plates and grown at 30^·^C for 3 days. Each strain carries both a [*URA3*] and a [*TRP1*] plasmid. The plasmids are either empty (EV) or carry the indicated *wtf25* alleles under the control of galactose-inducible promoters. The horizontal breaks in the images are due to rearrangements of the images to facilitate easy comparison. All strains within a panel were grown on the same plates (i.e., one SC-TRP-URA or SC Gal-TRP-URA plate). **E.** Representative images of the same strains depicted in D were induced in galactose for 4 hours at 30^·^C to express the indicated Wtf25 proteins. The images are not at the same brightness and contrast settings to clearly show localization of tagged proteins. The arrows in the TL panel highlight vacuoles. Yellow arrows indicate endoplasmic reticulum-like localization. 25^P^ indicates Wtf25^poison^, 25^A^ indicates Wtf25^antidote^, TL indicates transmitted light, and the scale bar is 4 µm.

**S14 Figure.**
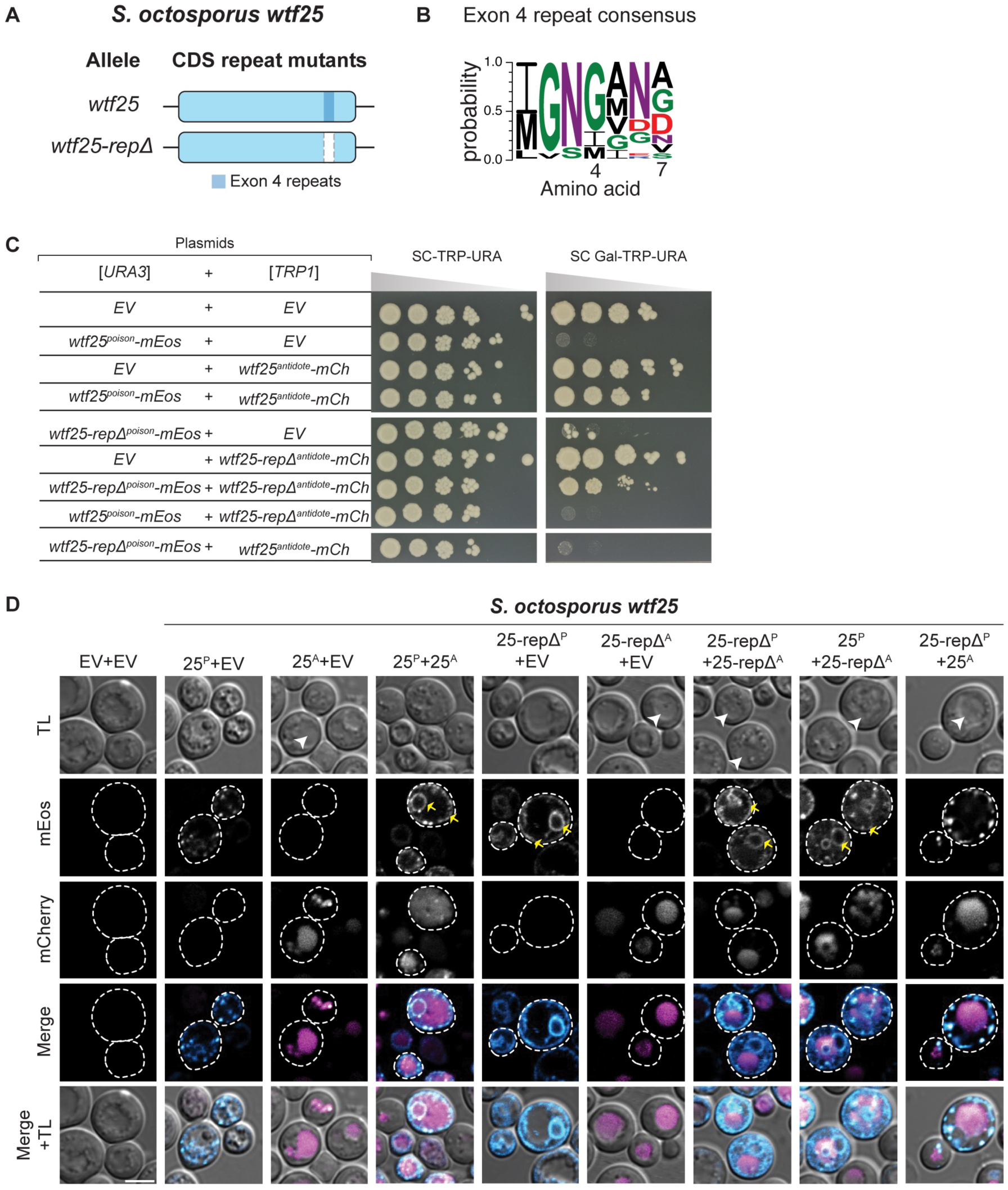
*S. octosporus wtf25* coding sequence repeats are functionally dispensable for poison toxicity but promote antidote rescue. **A.** Cartoon of *S. octosporus wtf25* exon 4 coding sequence repeat deletion mutant. **B.** Logo for the amino acids encoded by the repeats found in exon 4 of *S. octosporus wtf* genes from [19] **C.** Spot assay of cells serially diluted and plated on SC-TRP-URA and SC Gal-TRP-URA plates and grown at 30^·^C for 4 days. Each strain carries both a [*URA3*] and a [*TRP1*] plasmid. The plasmids are either empty (EV) or carry the indicated *wtf25* alleles under the control of galactose-inducible promoters. The horizontal breaks in the images are due to rearrangements of the images to facilitate easy comparison. All strains within a panel were grown on the same plates (i.e., one SC-TRP-URA or SC Gal-TRP-URA plate). **D.** Representative images the same strains depicted in C were induced in galactose for 4 hours at 30^·^C to express the indicated Wtf25 proteins. The images are not at the same brightness and contrast settings to clearly show localization of tagged proteins. The arrows in the TL panels highlight vacuoles. Yellow arrows indicate endoplasmic reticulum-like localization. 25^P^ indicates Wtf25^poison^, 25^A^ indicates Wtf25^antidote^, TL indicates transmitted light, and the scale bar is 4 µm.

**S1 Table. Overview of *wtf* alleles and their phenotypes in this study**. Columns 1-2 describe the allele names and their construction. Columns 3-4 describe the toxicity of the poison and antidote proteins encoded by the corresponding allele. Column 5 notes if the mutant antidote can rescue the corresponding poison allele. Column 6-7 note if the specific antidote allele can rescue the wild-type poison (Column 6), or if the specific poison can be rescued by the wild-type antidote (Column 7). Column 8 notes if the corresponding antidote can rescue other poison alleles. Column 9 notes the AmFRET values are comparable to the wild-type Wtf4^poison^-mEos or monomer-mEos. Finally, column 10 notes the figures/supplemental figures where these alleles can be found. Rows are separated based on the wild-type allele that the mutants were constructed from, where applicable. ND indicates that the specified experiment was not done.

**S2 Table. Overview of features across *wtf* driver genes.** Column 1 denotes the *wtf* driver gene, and columns 2-8 describe specific features of each gene and their encoded proteins. Column 2 notes if the gene has been shown to drive in fission yeast or been shown to encode for functional Wtf poison and antidote proteins in *S. cerevisiae*. Column 3 notes the number of exons in the gene, and column 4 notes the size of the Wtf^antidote^ protein in amino acids. If the antidote sequence had a predicted coiled-coil domain, column 5 notes the size and location in the antidote protein as described in [15]. Column 6 notes the numbers of PY motifs (L/PPXY; [24]) in exon 1. While some Wtf^antidote^ proteins also have an additional PY motif in exon 2, we did not include that information here since those motifs are shared by the corresponding poison proteins as well. Column 7 notes the number of predicted transmembrane domains predicted by TMHMM2.0 [72,73]. Column 8 notes the length of the exon 3 coding sequence repeats (if present) in amino acids. Column 9 notes the length of the exon 4/6 coding sequence repeats (if present) in amino acids. NA (not applicable) here indicates that the repeats were not present in the Wtf protein. Column 10 mentions the reference(s) where these genes/encoded proteins were characterized.

**S3 Table. Overview of predicted transmembrane domains of Wtf^antidote^ proteins.** The results from the TMHMM2.0 [72,73] analysis of Wtf^antidote^ proteins encoded by *wtf* driver genes in S2 table are detailed here. Find the specific results of the analyses, including the number of amino acids in the predicted transmembrane domains, probability of the N-terminal being internal to the membrane (N-in), the specific location and the length of the predicted transmembrane domains.

**S4 Table. Yeast strains used in this study.** Column 1 is the figure where these strains were used, Column 2 is the name of the yeast strain and column 3 is the genotype of the strain. If the strain was constructed in this study, column 4 has the details on how the strain was constructed. If the strain was constructed in another study, column 5 notes the references for the same.

**S5 Table. Plasmids used in this study.** Column 1 denotes if the plasmids were used for strain construction or cloning, and the specific wild-type alleles that the plasmids correspond to. Column 2 is the name of the plasmid and column 3 is a short description of the plasmid. If the plasmid was not constructed in this study, column 4 references the studies the plasmids were constructed in.

**S6 Table. Oligos used in this study.** Column 1 denotes the name of the oligo and column 2 has the sequence of the oligos. If the oligo was not constructed in this study, column 3 references the studies the oligos were used in.

**S1 Data. Raw Data for Pearson’s Correlation tests performed in this study.** Each sheet contains the raw data for the Pearson’s Correlation tests and the statistical analysis corresponding to the tests performed in Figs 4F, S6E, 6F and S11F, respectively. See Methods for how this analysis was performed.

**S2 Data. Raw Data for AmFRET plots presented in this study.** Each sheet contains the raw data and the statistical analysis for the AmFRET plots of different Wtf^poison^-mEos proteins, Wtf4^antidote^-mEos, *S. kambucha wtf4* Exon1-mEos and monomer-mEos proteins in Figs 2B, S2E, S3D, Fig 4C, S6 Fig, S7D and S8B, respectively. The corresponding raw data for the Acceptor fluorescence intensity presented in S4 Fig for each of these experiments are found next to the relevant AmFRET data. See Methods for how this analysis was performed.

